# Chemical inhibition of histidine biosynthesis curtails *M. tuberculosis* infection

**DOI:** 10.1101/2023.06.20.545697

**Authors:** Satish Tiwari, Mohammed Ahmad, Varun Kumar, Deepsikha Kar, Swati Kumari, Abhisek Dwivedy, Ravi Kant Pal, Amit Kumar Mohapatra, Vaibhav Kumar Nain, Vishawjeet Barik, Rahul Pal, Ranjan Kumar Nanda, Amit Kumar Pandey, Bichitra Kumar Biswal

**Author notes:** Department of Bioengineering, University of Illinois, Urbana Champaign (Abhisek Dwivedy).

## Abstract

To overcome the drug resistance crisis and shorten the current duration of human tuberculosis (TB) therapy, new anti-TB molecules is required. In an earlier study, we have shown that *Mycobacterium tuberculosis* (*Mtb*), the causative agent of TB, with a fractured de novo histidine biosynthesis fails to mount TB infection in mouse model, emboldening that disrupting the function of this pathway may constitute a novel strategy to curtailing TB infection. In this study, through a target based approach we have designed a number of triazole scaffold molecules specific to imidazole glycerol phosphate dehydratase (IGPD; HisB) of this pathway and have delineated atomic level interactions between the enzyme and inhibitors which pinpointed the specificity and the inhibitory mechanism. Importantly, these molecules exhibited significant potency against free as well as macrophage-internalized wild-type and drug-resistant clinical isolates in culture medium. Notably, a couple of these compounds showed efficacy in reducing the bacterial burden in *Mtb*-infected mouse model. The chemical inhibition of IGPD induces histidine auxotrophy in *Mtb* and brings in new prospects to the area of anti-TB drug discovery.

## Introduction

Human tuberculosis (TB), caused by the bacterial pathogen *Mycobacterium tuberculosis* (*Mtb*) continues to be a major health issue worldwide^1^. The antibiotic resistance by *Mtb* poses major challenges in effectively clearing the infection by the currently available anti-TB therapeutics^2^. The major targets of the current drugs are the cell wall and protein biosynthesis^3^, however the bacillus has developed significant resistance to drugs that target these biological processes. Even the occurrence of drug resistance against bedaquiline, recently developed drug to treat drug resistance TB, has been documented^4, 5^. Negative side effects associated with bedaquiline treatment adds another layer to the problems^6^. To reduce current anti-TB treatment time, abate drug resistance crisis and combat this ever evolving human pathogen, exploring new options that could limit *Mtb* infection is certainly required. In the past decades, a vast amount of progress made in studying the TB biology has assisted in identifying a plethora of essential factors that the bacillus employs to evade host strategies directed in its clearance, enabling it to grow in an adverse host environment. Among various such factors, amino acids biosynthesis by *Mtb* play major roles in mounting sustained TB infections^7–10^. For example, a host strategy, to kill the pathogen, starves *Mtb* of Histidine (His); however *Mtb* manages its His requirement through the de novo histidine biosynthesis pathway^8^. This suggests that disrupting the function of this pathway may curtail bacterial growth and therefore may represent a new way of limiting TB infection. Furthermore, the fundamental requirement of His for *Mtb* viability and the inability of humans to not make His de novo^11^ make this pathway an attractive drug target. In a major focus to design new anti-TB small molecules with novel mechanism of action, over the years, we have structurally and biochemically characterized IGPD (also known as HisB)^12, 13^, HisC^14^ and HisN^15^ of this pathway. Notably, primarily owing to the absence of its both structural and functional homologs in mammals, IGPD belongs to a prime category of anti-bacterial^13^, anti-herbs^16^ and anti-fungal^17^ drug target. In this study, through a target-based approach, we have identified a number of anti-HisB inhibitors and show that chemical inhibition of the *Mtb* de novo histidine biosynthesis leads to a significant bacterial clearance. Taken together, this study identifies new chemical entities as anti-TB compounds specific to *Mtb* IGPD which in turn implicate in expanding the current anti-TB lead molecules library.

## Results

### Identification of Triazole scaffold molecules as HisB inhibitors

Over the last 25 years, plant IGPD, an enzyme that catalyzes the sixth step of the His pathway (Fig S1A), has been an important herbicide target^16^. A plethora of studies particularly in the area of developing potent herbicides targeting IGPD have led to the discovery of a ample number of small molecules^16, 18, 19^. These studies showed that triazole and imidazole scaffold compounds are promising IGPD inhibitors. The availability of such a wealth of information provides a basis to examine whether triazole and imidazole scaffold inhibitors exhibit anti-TB potency by inhibiting the function of the His pathway of *Mtb.* Our earlier studies^12, 13^ and reports from other groups^16, 18^ suggest that triazole and imidazole derivatives compounds have better prospects in inhibiting the function of IGPD. The 3D experimental structure of this enzyme from *Mtb* has been elucidated by us^12^ and from other organisms by others^20, 21^. The results from our study and those of others show that the functional unit of the enzyme is a 24-mer (Fig S1B) with 24 identical active sites. Each active site is asymmetric and made up with three subunits (Fig S1C and S1D). By utilizing these structural information into the inhibitors design; we constructed a number of triazole derivative compounds (Table S1) and calculated the binding affinity of each of these compounds. The physicochemical properties together with the binding scores (Table 1A and Table 4) clearly suggest that these molecules have anti-TB potential and therefore warranted further investigation. In order to pursue further, we explored the commercial availability of these molecules and a few of these molecules [we named them –SF1, SF2 and SF3 (Fig 1A)] were procured from Sigma chemicals, USA and their in vitro and in vivo specificities were investigated.

**Figure 1.**
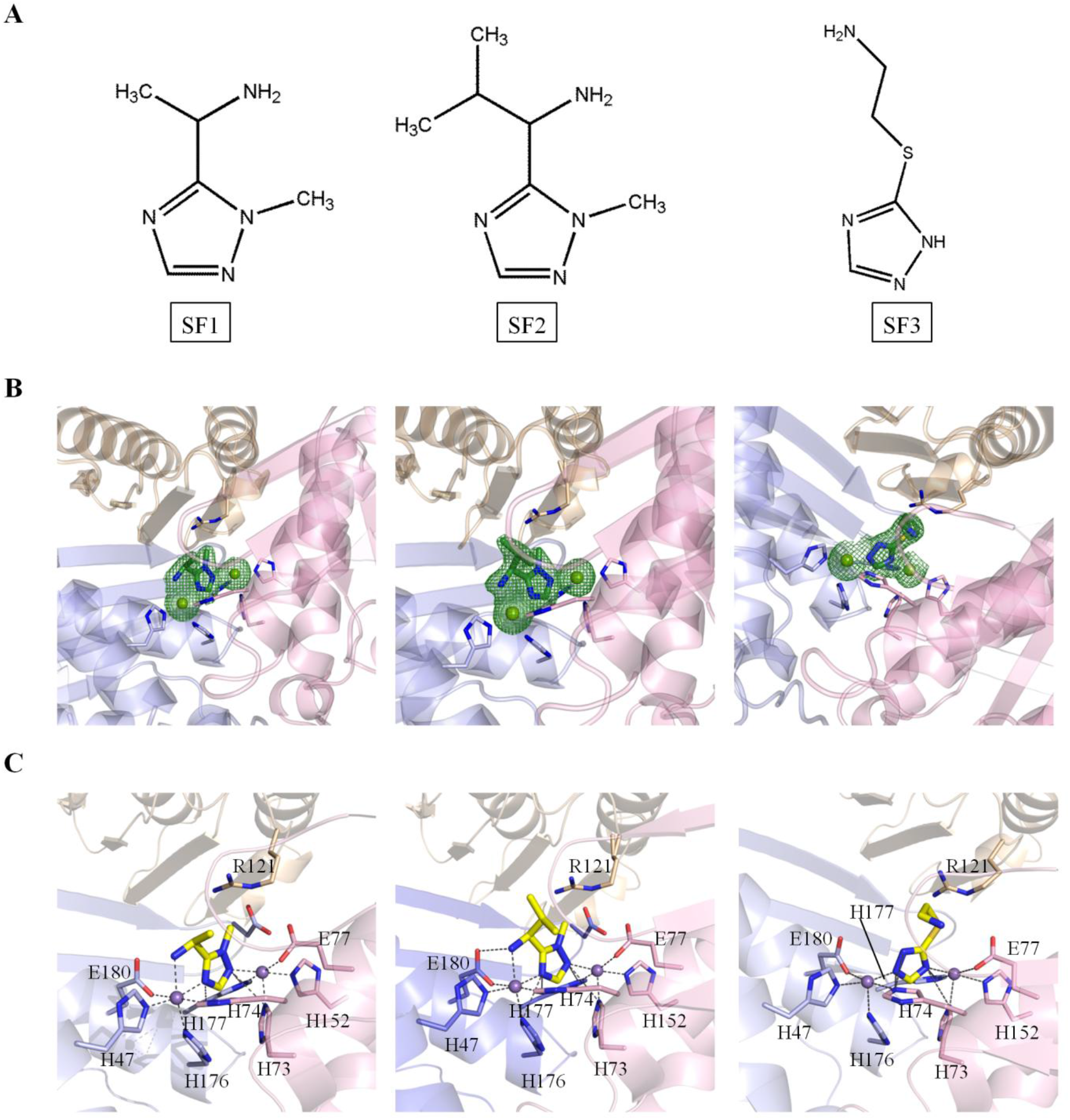
General Structures, Binding and HisB/inhibitors interactions. (A) Chemical structure of the inhibitors SF1, SF2, and SF3. (B) Electron density maps of enzyme/inhibitor complexes. The difference Fourier map *|F_o_|* - *|F_c_|* (shown in green grids) at 3σ contour level for all three inhibitors suggest that these inhibitors bind at the active site pocket of the enzyme. (C) Interactions (shown in black dashes) of HisB with SF1, SF2, and SF3 inhibitors. The amino acid residues involved in the interactions are shown in stick representation, and two Mn^2+^ are shown in spheres with the Purple-blue color.

**Table 1:**
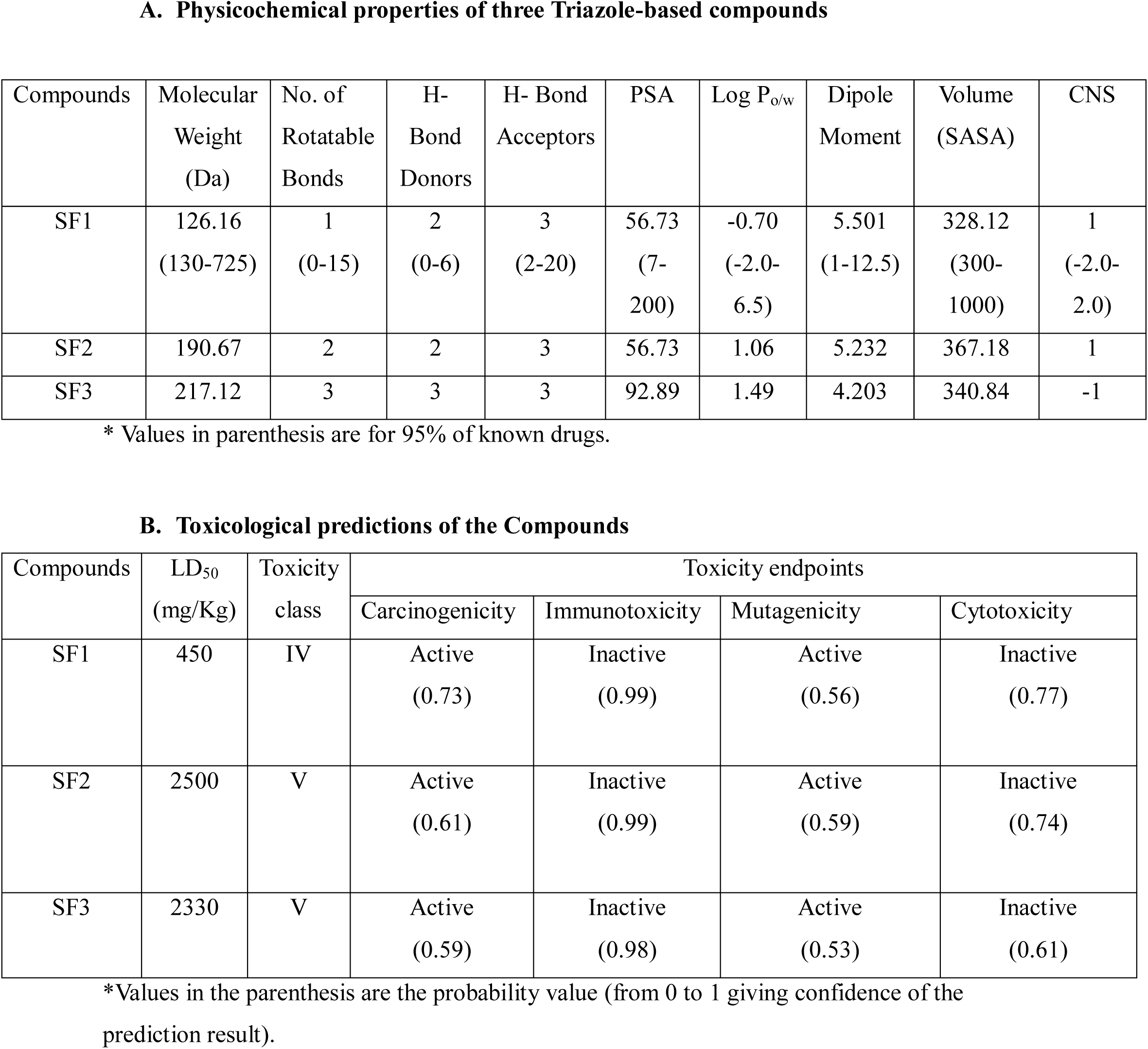

A. Physicochemical properties of three Triazole-based compounds
| Compounds | Molecular Weight (Da) | No. of Rotatable Bonds | H- Bond Donors | H- Bond Acceptors | PSA | Log P<sub>o/w</sub> | Dipole Moment | Volume (SASA) | CNS |
| --- | --- | --- | --- | --- | --- | --- | --- | --- | --- |
| SF1 | 126.16<br>(130-725) | 1<br>(0-15) | 2<br>(0-6) | 3<br>(2-20) | 56.73<br>(7-200) | -0.70<br>(-2.0-6.5) | 5.501<br>(1-12.5) | 328.12<br>(300-1000) | 1<br>(-2.0-2.0) |
| SF2 | 190.67 | 2 | 2 | 3 | 56.73 | 1.06 | 5.232 | 367.18 | 1 |
| SF3 | 217.12 | 3 | 3 | 3 | 92.89 | 1.49 | 4.203 | 340.84 | -1 |

\* Values in parenthesis are for 95% of known drugs.

B. Toxicological predictions of the Compounds
| Compounds | LD<sub>50</sub><br>(mg/Kg) | Toxicity class | Toxicity endpoints |  |  |  |
| --- | --- | --- | --- | --- | --- | --- |
|  |  |  | Carcinogenicity | Immunotoxicity | Mutagenicity | Cytotoxicity |
| SF1 | 450 | IV | Active<br>(0.73) | Inactive<br>(0.99) | Active<br>(0.56) | Inactive<br>(0.77) |
| SF2 | 2500 | V | Active<br>(0.61) | Inactive<br>(0.99) | Active<br>(0.59) | Inactive<br>(0.74) |
| SF3 | 2330 | V | Active<br>(0.59) | Inactive<br>(0.98) | Active<br>(0.53) | Inactive<br>(0.61) |

\*Values in the parenthesis are the probability value (from 0 to 1 giving confidence of the prediction result).

### Triazole scaffold compounds show drug like properties

In order to determine their suitability as potential anti-TB compounds, we analyzed the physicochemical properties of the aforementioned compounds. These molecules show appropriate gastrointestinal absorption and low penetration of the blood-brain barrier similar to that of one of the first-line TB drugs, isoniazid (INH). Qikprop analysis categorized these molecules as drug-like with zero violation of a widely regarded, “Lipinski rule of five” (Table 1A), and most of the properties of the compound were in the range of 95% known drugs.

ProTox Cytotoxicity predictions suggested mild probabilities for compounds to be toxic or carcinogenic (Table 1B).

### Determining Molecular level specificities by X-ray crystallography

Elucidating enzyme/inhibitor interactions greatly assists in understanding the mechanism of drug action and helps in improving molecular design to achieve higher potency. In this regard, we determined the crystal structures of IGPD-inhibitor complexes individually. A search for non-protein electron density peaks, particularly in the active site area, showed the presence of significant 2|*F_o_*| – |*F_c_*| and |*F_o_*| – |*F_c_*| peaks resembling the shape of these inhibitors, demonstrating the inhibitors’ binding at the active site of the enzyme (Fig 1B). The critical requirements for the binding at the active site are a ring which allows the formation of coordination bonds to the Mn^2+^ ions and a negatively charged group separated by 2-3 carbon chain from the ring. The triazole ring of each inhibitor is anchored between these two active site manganese ions (Fig 1C). The binding mode of the triazole ring is the same for all inhibitors (Fig S2). The two active site manganese atoms are coordinated with the two nitrogen atoms of the triazole ring. The active site is made up of mainly histidine-rich amino acids.

### Characterization of the kinetics parameters

In order to examine the degree of affinity and specificity of each enzyme/inhibitor complex, we carried out in vitro kinetics analysis using a protocol reported by us in an earlier study^22^. The IC_50_ of inhibitors lie in the range of 33.46 µM (SF1), 44.09 μM (SF2), and 76.95 μM (SF3), calculated by plotting log (inhibitor concentration) vs response, four-parameter variable slope curve (Fig 2A). These compounds inhibit the growth of H37Rv as well as drug resistant clinical isolates (S6 –Resistant to INH+Rifampicin & S7 –Resistant to INH) in minimal media with MIC_99_ values in micromolar range (Table 2) as confirmed by alamarBlue assay (Fig 2B). Further, we examined the effect on the growth of *Mtb* inhibited by these compounds following histidine supplementation in minimal media. As expected, histidine supplementation nullifies the inhibitory effects of the compounds and restores the *Mtb* growth (Fig 2C), confirming that these compounds indeed target the histidine biosynthesis.

**Figure 2.**
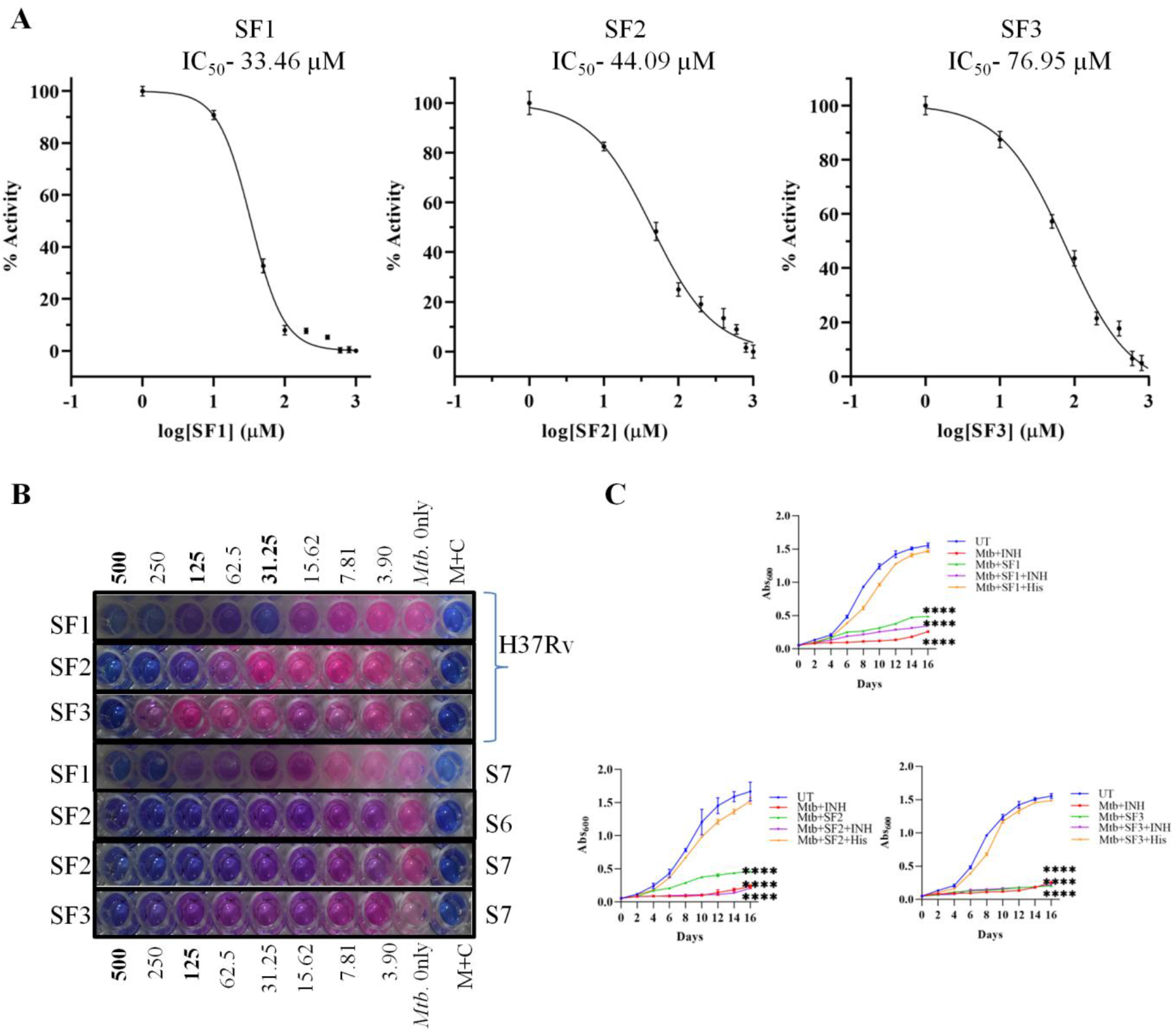
Effect of Triazole based compounds SF1, SF2, and SF3 on *Mtb* growth in vitro. (A) Activity of compound against the enzyme IGPD; HisB for IC_50_ values. All the Data are shown as biological duplicates with mean ± s.d (n=2). (B) MIC_99_ values against H37Rv and drug resistant clinical isolates (S6 & S7) determined by alamarBlue assay. All the experiments are done at least in biological duplicates. [Media + Compound = M+C] (C) Growth curve analysis showing the effect of SF1, SF2, and SF3 on *Mtb* growth at a concentration of 1x MIC for 14 days with and without histidine. All the data points are mean ± s.d of three independently grown bacterial cultures (n=3).

**Table 2:** Potency (MIC_99_ values) of compounds against H37Rv and drug resistant clinical isolates.

| Compounds | MIC<sub>99</sub> (H37Rv) μM | MIC<sub>99</sub> (S6) μM | MIC<sub>99</sub> (S7) μM |
| --- | --- | --- | --- |
| SF1 | 31.25 | - | 250 |
| SF2 | 125 | 125 | 125 |
| SF3 | 500 | - | 250 |

### Triazole compounds clear Mtb infection in macrophages

To examine the in vitro potency of these compounds, we infected macrophages cell lines (RAW 264.7) with *Mtb* at Multiplicity of Index (MOI) = 10 and observed the bacterial load in the presence of various concentrations of each compound. As shown in Fig 3, bacterial load is reduced significantly after 3 days of treatment, showing approximately a thousand-fold decline for SF1 (Fig 3A) and hundred-fold decline for SF2 treatment (Fig 3B). Although, SF3 compound inhibited the IGPD activity and curbed *Mtb* growth in culture medium but did not reduce the bacterial load in *Mtb* infected– macrophages at concentrations up to 1x of MIC_99_.

**Figure 3.**
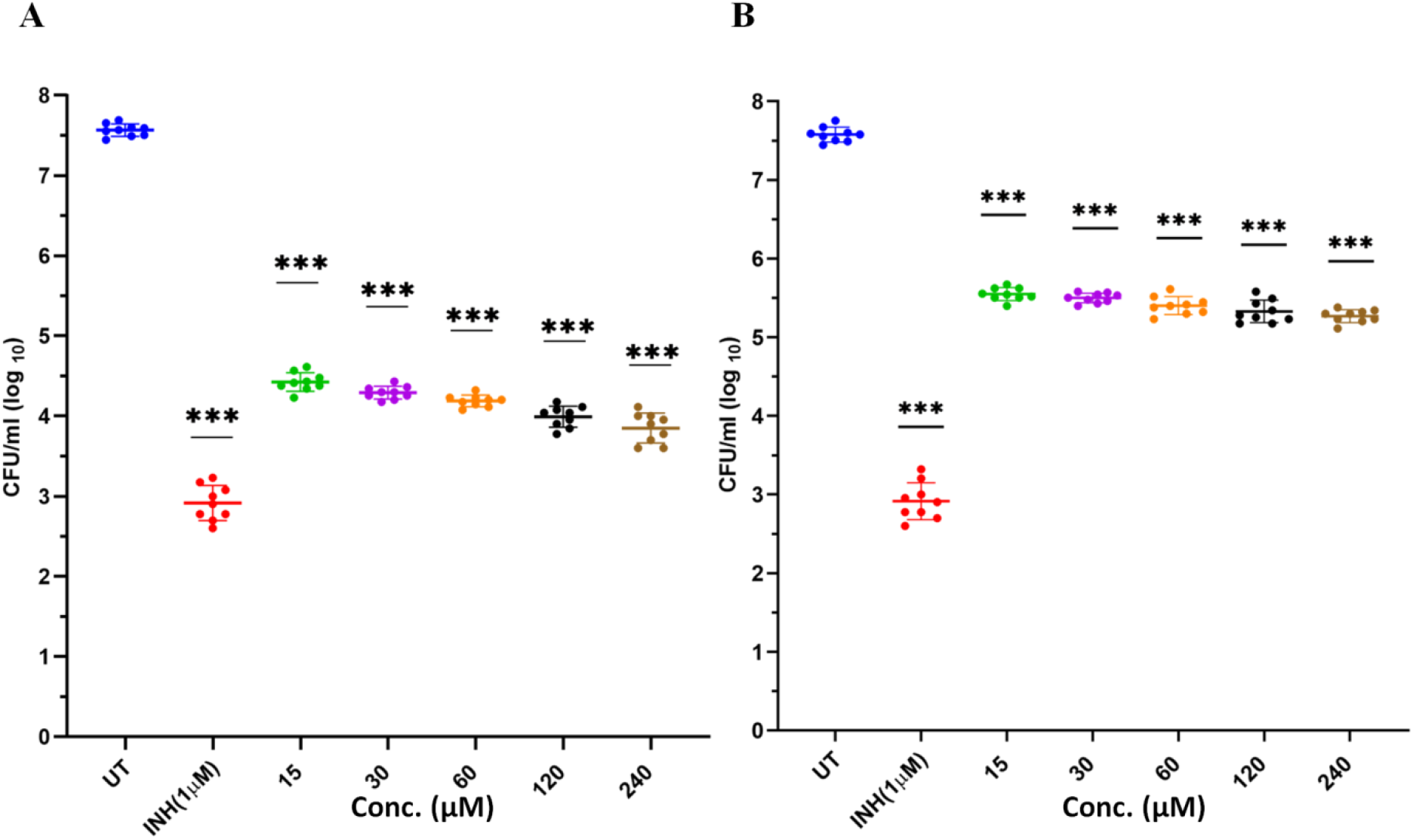
Efficacy testing in *Mtb* infected macrophage model. The intracellular killing efficacy of Triazole based compounds in *Mtb* infected murine macrophage (RAW264.7) model. RAW264.7 macrophages were infected with H37Rv at an MOI of 10 for 3 h before treatment with different concentrations of compounds (15, 30, 60, 120, and 240 μM), in comparison with INH (1 μM). (A) Effect of SF1 compound on *Mtb* after treatment for 72 h is represented by log CFU/ml. (B) Effect of SF2 compound on *Mtb* after treatment for 72 h is represented by log CFU/ml. One-way Anova test (Dunnett’s multiple comparison test) was used to calculate P-value (\**\*\*p<0.001)*. Data are shown as mean ± s.d for 3 biological replicates (n = 3).

### Evaluation of in vitro and in vivo toxicity of these compounds

The water-soluble properties of these inhibitors make them better prospects for absorption. We tested their toxicity both at in vitro and in vivo systems. Using alamarBlue assays in Raw 264.7, THP1 and HEK-293 cell lines; we observed that these compounds are non-toxic even at millimolar range concentrations (Fig S3). We further treated mice with different doses of each of these compounds and observed that all mice were alive after an oral administration of a single dose of 5, 50, 500, and 2000 mg/kg body weight of SF2 compound, while 2 mice of 2000 mg/kg body weight and 3 mice of 5000 mg/kg body weight died after administration of SF3 compound within four days (Table 3A and B). This suggested that the LD_50_ value for SF2 is greater than 2000 mg/kg body weight and that of SF3 is below 5000 mg/kg body weight. All surviving mice behaved normally with no clinical signs of adverse effects / toxicity caused by the compounds after oral administration of the dose of 5, 50, 500 and 2000 mg/kg body weight of SF2 and SF3 during the entire 14 days acute oral toxicity study. Clinical signs such as slowness and weight decline appeared after 12-24 h in mice, after administration of single dose of 5000 mg/kg of SF3. After single dose of oral administration, mice body weight generally increased for all dose groups throughout the study (Fig S6). Compared to the untreated group, body weight of mice administered with SF2 increased for all doses while a slight dip in weight for 5 mg/kg body weight on 7^th^ day was observed. For SF3 treatments, body weight decreased for 5, 500, and 2000 mg/kg body weight and increased for 5000 mg/kg body weight compared to the untreated group was seen.

**Table 3:** Analysis of the acute oral toxicity test of SF2 and SF3 on CD1 mice in 14 Days study. Doses (mg/kg body weight) and Number of deaths (D).

**A.**
| Compound | Doses (mg/kg) | Mice number initial | Mice number final | Total Deaths (D) |
| --- | --- | --- | --- | --- |
| SF2 | 5 | 5 | 5 | 0 |
|  | 50 | 5 | 5 | 0 |
|  | 500 | 5 | 5 | 0 |
|  | 2000 | 5 | 5 | 0 |

**B.**
| Compound | Doses (mg/kg) | Mice number initial | Mice number final | Total Deaths (D) |
| --- | --- | --- | --- | --- |
| SF3 | 5 | 5 | 5 | 0 |
|  | 50 | 5 | 5 | 0 |
|  | 500 | 5 | 5 | 0 |
|  | 2000 | 5 | 3 | 2 |
|  | 5000 | 5 | 2 | 3 |

### Triazole scaffold compound show non-toxic effects in mouse model

All the biochemical analysis was done using semi-automated machine, Coralyzer. After performing the serum analysis of SF2 and SF3 treatments for all doses, no significant changes were observed in the levels of Serum Glutamic Oxaloacetic Transaminase (SGOT), Serum Glutamic Pyruvic Transaminase (SGPT), creatinine, and urea, compared to the untreated group. As all values for the hepatic function marker enzymes (SGPT, SGOT) and renal and hepatic function marker enzymes (urea, creatinine) were in the normal range, it indicated no toxicity to liver and kidney (Fig 4).

**Figure 4.**
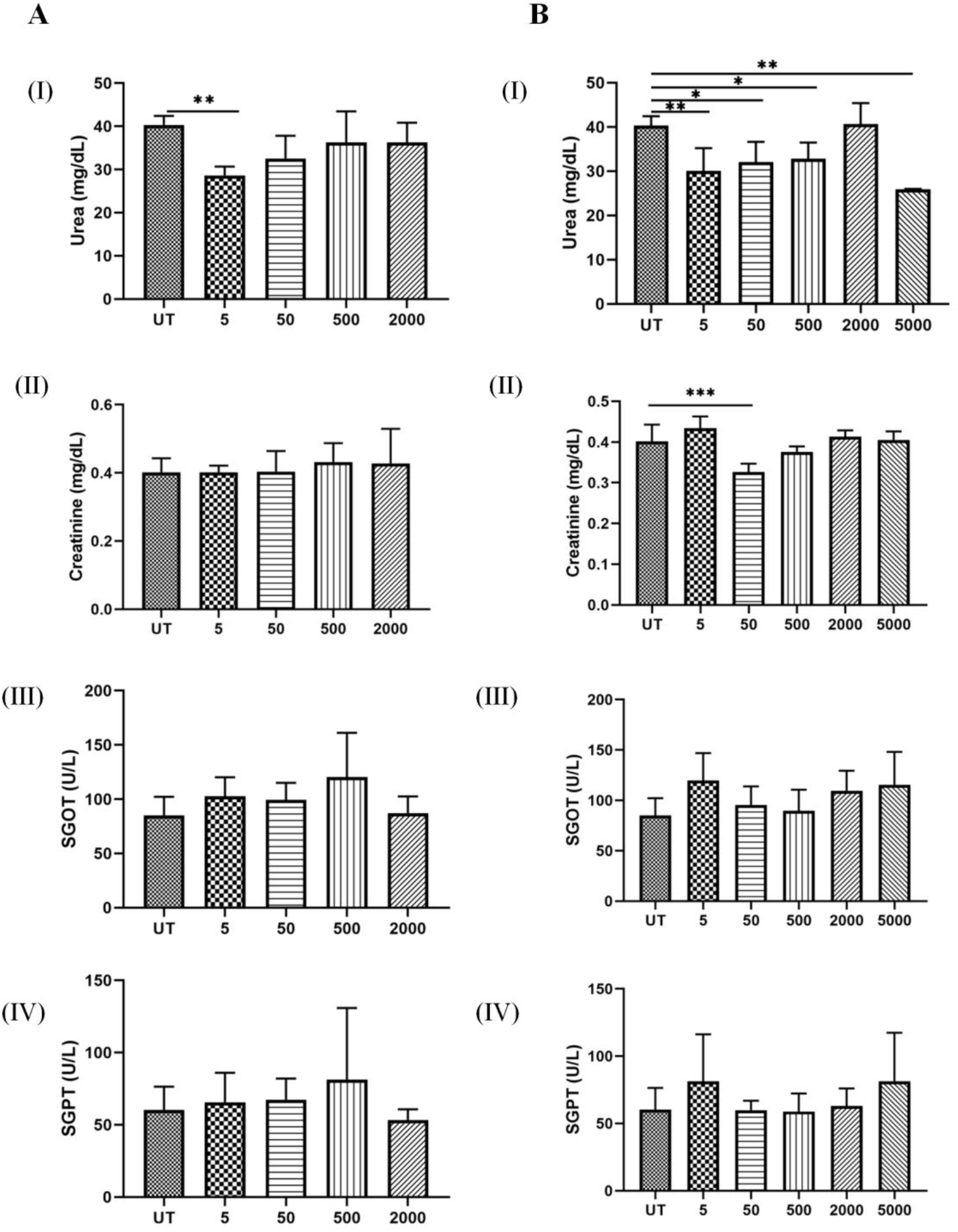
Animal toxicity data showing biochemical parameters. Effect of varying doses of SF2 and SF3 compound on (I) Urea, (II) Creatinine, (III) SGOT, and (IV) SGPT in different groups. (A) Untreated group and 4 treated groups (administered with - 5, 50, 500, and 2000 mg/kg body weight of SF2) [n=5]. (B) Untreated group and 5 treated groups (administered with - 5, 50, 500, 2000 and 5000 mg/kg body weight of SF3) [n=5].

Further, we performed the histopathological analysis to look for the tissue toxicity caused by the compounds used in our study. No significant changes were observed in the kidney and liver sections for the compounds SF2 and SF3. Histopathological parameters of liver and kidney in treated and the untreated groups were observed and confirmed by H&E staining. Histopathological examination of kidney with normal renal cortex, no leukocyte infiltration, no signs of edema, absence of necrotic foci, and normal tubular morphology while same stands for liver with normal Kupffer cells and no signs of granuloma, necrosis and infiltrations following SF2 treatment. Similarly, SF3 treatment showed no changes in the gross kidney histopathology at different doses (Fig S4 and S5).

### Testing the efficacies of these compounds in Mouse model of TB infection

Based on absorption, cytotoxicity and in vitro macrophage clearance data, these inhibitor molecules were further tested for their efficacy against *Mtb* in animal model of TB infection. Briefly, *Mtb* infected BALB/c mice were administered with test compounds SF1 (100 mg/kg body weight), SF2 (100 mg/kg body weight), SF2 (2x) (200 mg/kg body weight) and SF3 (100 mg/kg body weight) 4 weeks post-infection (Fig 5A). Eight weeks post-infection the mice were euthanized and the bacterial burden of the infected lungs and spleen were estimated by performing a colony forming unit assay (Fig 5B and C).It was observed that in comparison to the untreated control group, the lungs and spleen of mice treated with test compounds SF1and SF2 showed no reduction in the bacterial burden (Fig 5D and F). However, compound SF3 at the same concentration demonstrated significant lowering of the bacterial burden in both lungs and spleen relative to the untreated (UT) mice, (Fig 5E and G). We also tested the SF2 compound at twice the concentration and observed that in comparison to the UT control group; there was a significant decrease in the bacterial load in the lungs of the mice (Fig 5E). However, the reduction in the bacterial load in the infected spleen was not found to be statistically significant, (Fig 5G). These above findings indicate that the compound-mediated histidine limitation compromised bacterial survival inside the host microenvironment. Interestingly, supplementation of histidine in mice treated with SF2 (2X) and SF3 restored the bacterial burden to the untreated levels further validating the fact that the decrease in the bacterial load observed in the treated mice was a result of reduced biosynthesis of histidine.

**Figure 5.**
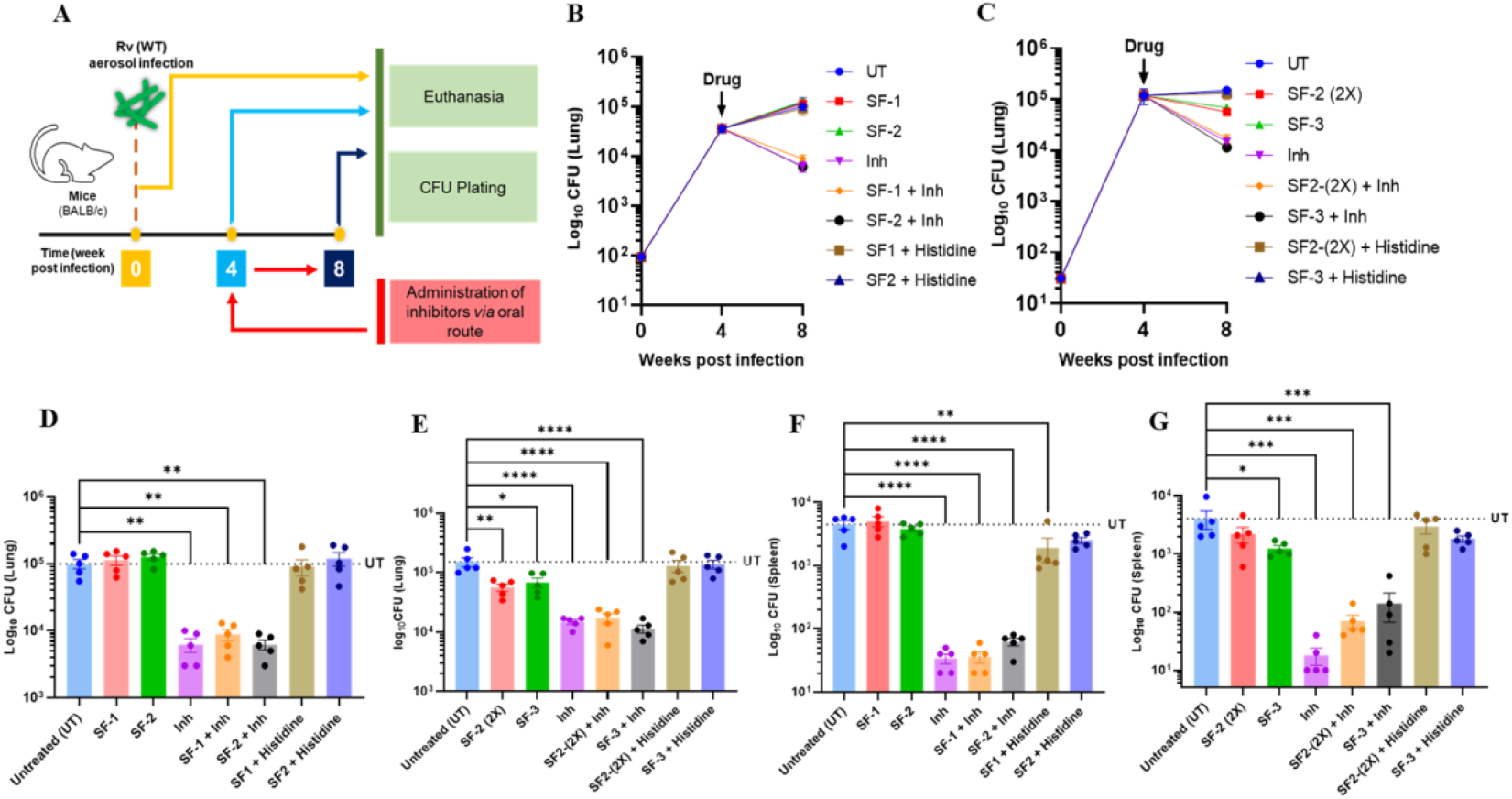
In-vivo efficacy. Effect of test compounds - SF1, SF2 (1x), SF2 (2x) and SF3 on clearance of *Mtb* in mice was determined as depicted in schematic (A). Mice lungs and spleen bacterial burden were evaluated post administration of compounds orally at indicated time points (B and C). Clearance by SF1 and SF2 was comparable to the bacterial load in untreated group (UT) in both lungs and spleen (D and F). SF2 (2x) and SF3 showed significantly reduced bacterial burden than control (UT) group in lungs (E). Similarly, SF3 demonstrated significantly lower bacterial count in spleen, so was in case of SF2 (2x), albeit not significant (G). The CFU counts amongst the tested groups were analysed by One-way ANOVA (Sidak’s Multiple Comparison Test) with *\*\*\*\*p<0.0001, ***p = 0.0001, **p<0.001, *p<0.01*.

## Discussion

To shorten the current anti-TB drug regimen and overcome the antibiotic resistance crisis, the future of TB treatment requires novel therapeutic molecules with established mechanisms of action. The discovery of bedaquiline^23^ and delamanid^24^ both to treat drug resistant TB almost after over 40 years since the launch of the first-line anti-TB drugs have emboldened TB drug discovery researchers. Unfortunately, the bacillus has developed resistance against these drugs^4^. Moreover the use of bedaquiline has been associated with serious side effects^6^. Although the phenotypic approach has majorly contributed to the discovery of anti-TB drugs, the target based approach has recently gained momentum. Owing to the fundamental requirement of amino acids in many critical cellular processes, the major one being protein synthesis, disrupting their biosynthesis could lay new paths in combating TB. The study on His pathway has revealed that disrupting the function of this pathway may yield a new generation of anti-TB lead molecules. Target based approach has helped to identify compounds that showed target activity in cell-free medium and whole-cell systems^25^. These results show that deriving detailed information about the criticality of a target could potentially assist to identify novel anti-TB molecules. Targeting *Mtb* His pathway is particularly advantageous for two reasons-(1) the pathway is critical for *Mtb* to mount a sustained TB infection in vivo and (2) it is absent in humans, meaning pathway enzymes lack human homologs. This implies that developing inhibitors of these enzymes would minimize the potential of cross-reactivity. IGPD being an established anti-herb target for past 25 years, the vast amount of available data assisted us to figure out probable initial candidate molecules. Our earlier studies on deciphering the interactions between IGPD and the substrate [Imidazole glycerol phosphate (IGP)] combined with existing knowledge of active compounds laid a profound platform in the design of molecules. In this study we employed a rational approach to design new IGPD inhibitors disrupting bacterial His pathway. Our findings will assist in designing new analogues of these compounds with improved efficacy and specificity by employing a structure-activity relationship strategy.

Toxicological studies have shown that in acute oral toxicity tests, SF2 and SF3 do not produce any toxic/adverse effects in mice. Furthermore, SF2 caused no mortality or morbidity in any mice that survived the 14 days of observation. The LD_50_ value of SF2 was greater than 2000 mg/kg body weight. Based on the present study, we have predicted the dose of the compound at which it is safe and non-toxic after oral administration. In addition to this, there were no significant changes in biochemical parameters (SGPT, SGOT, urea, and creatinine) in any of the studied groups treated with a single dose of SF2 and SF3, which are usually measured to assess any toxic effects of compounds on the liver and kidney. The liver and kidneys were analyzed for histopathological changes such as leukocyte infiltration, necrosis, edema and vacuoles. These histopathological examinations are the gold standard for assessing the post-treatment related pathological effects in tissues and organs. All assessed organs were found to be normal on histopathological examination, with no signs of toxicity in any of the tested groups with respect to biochemical parameters, body weight and histopathological examination. Finally, it can be concluded that SF2 and SF3 are non-toxic, with no observed adverse effect level (NOAEL) greater than 2000 mg/kg body weight for SF2 and less than 5000 mg/kg body weight for SF3. Finally, compounds SF2 and SF3 cleared TB in a mouse model of infection at dosage concentrations as low as 100 and 200 mg/kg body weight, respectively, further highlighting the anti-tubercular potential of these compounds in an actual clinical setting. Taken together, this study identified HisB-specific anti-TB molecules that act by abolishing the function of an essential metabolic pathway in pathogens, representing a novel way of curtailing TB infection.

## Conclusion

We have demonstrated, in an earlier study, that the de novo histidine biosynthesis is required for *Mtb* to escape the host immune mediated clearance^8^, quelling a common notion that *Mtb* meets its histidine requirement by importing from the host cellular milieu. This feature emphasizes the significance of targeting histidine biosynthesis pathway for designing/developing novel anti-TB molecules. In this study, through a target-based approach, we have identified a number of triazole-based compounds specific to the HisB. The crystal structures of HisB/inhibitor complexes aided in mapping the interactions between the enzyme (IGPD; HisB) and inhibitors (SF1, SF2, and SF3). We also show that these compounds are non-toxic at in vitro as well as in vivo conditions, making these molecules attractive anti-TB drug candidate. More importantly, these compounds are able to clear bacterial load in mouse model of TB infection. Collectively, this study has brought up new insights in the area of anti-TB drug discovery.

## Experimental Section

### Making up small molecule library and screening on IGPD

One active site of IGPD consisting of three subunits was constructed and the active site area was chosen as a grid to dock each small molecule in this area. Features of small molecules, such as their structure and shape, play an important role in the interaction of residues present in the active site of the protein. This information could help build a specific and influential library of small molecules. Over the years, many compounds have been researched to identify drugs against TB, among which triazoles have been found to be the most explored and valuable scaffold. Triazole is a heterocyclic compound that includes a five-membered ring consisting of two carbon and three nitrogen atoms. The properties of triazole rings, such as hydrogen-binding ability, moderate dipole moment, and enhanced water solubility under in vivo conditions, are responsible for their enhanced biological activities. Ongoing research is being carried out across the globe to seek effective treatments for TB, which has suggested that a compound with a triazole motif is a promising inhibitor of IGPD^13, 26–28^. Using this as a basis, we created a library of compounds with triazole motifs. First, we converted the 3-Methyl-1H-1,2,4-triazole structure into a Simplified Molecular Input Line Entry System (SMILES), a 2D representation, and the chemical language of the chemical structure^29^. The SMILES string for 3-Methyl-1H-1,2,4-triazole is CC1=NC=NN1. Subsequently, the 2D representation was searched in the PubChem database^30^, where the closest structure to the triazole motif was identified and chosen for creating the triazole motif-specific in-house library. A total of 50 compounds were shortlisted and set up for in silico virtual screening of the prepared HisB receptor. In-house library compounds with their specific codes, SMILE string, and Molecular Formula are summarized in Table S1. The structures of the compounds are listed in Table S2. Each of these molecules was docked in this area using a guided docking protocol with Schrodinger drug discovery software^31–33^. The docking score and binding energy are listed in the Table 4. All the compounds used in the study were procured from Sigma, USA and are >95% pure as reported by Sigma.

**Table 4:** Docking scores and binding energies of top 15 compounds.

| Code | Docking score (kcal/mol) | MMGBSA dG Bind (kcal/mol) |
| --- | --- | --- |
| SF14 | -6.539 | -8.40 |
| SF18 | -5.701 | -18.38 |
| SF5 | -5.589 | -27.20 |
| **SF3** | **-5.587** | **-29.59** |
| SF44 | -5.546 | -28.09 |
| SF15 | -5.317 | -20.41 |
| SF34 | -5.184 | -9.66 |
| **SF1** | **-4.936** | **-11.97** |
| SF36 | -4.863 | -8.44 |
| SF37 | -4.713 | -34.68 |
| SF43 | -4.493 | 5.26 |
| SF42 | -4.435 | -26.53 |
| SF12 | -4.370 | -25.71 |
| **SF2** | **-4.368** | **-11.65** |
| SF13 | -4.327 | -20.11 |

### Over expression and purification of IGPD

The over-expression of the enzyme in a *Mycobacterium smegmatis (Msmeg)* over expression system and its purification to a high degree of homogeneity were carried out using protocols that we have established earlier^34^. Briefly, the Rv1601-pYUB1062 expression vector was electroporated into competent *Msmeg* strain mc^2^4517 cells. A single transformed colony was revived in 10 ml of Luria Bertani (LB) broth supplemented with 0.05% Tween 80, 0.2% glycerol, and the antibiotics kanamycin (25 µg/ml) and hygromycin B (50 µg/ml). The culture was grown at 310 K, and 180 rpm for 24 h. 1 ml of the primary culture was inoculated into 50 ml of the same medium and grown at 310 K, 180 rpm until A_600_ reached 0.6 – 0.8. Subsequently, the culture was diluted 30-fold into 1.5 liters of the same medium to make a secondary culture and grown to mid-exponential phase (A_600_ =0.8) at 310 K, 180 rpm and then induced with 0.02% acetamide. After 24 h of induction, cells were harvested by centrifugation at 10,000 x g for 20 min. The cell pellet was resuspended in 50 ml of 20 mM Tris, pH 8.5, 200 mM NaCl, 5% glycerol, and 20 mM imidazole buffer with one Complete Mini, EDTA-free protease inhibitor tablet (Roche Applied Science). The cells were lysed at 277 K under high pressure (25,000 p.s.i.) using a cell disrupter (Constant Systems Ltd., UK). The lysate was centrifuged at 10,000 x g for 45 min at 277 K to the remove unbroken cells and inclusion bodies. The supernatant was then loaded onto an equilibrated nickel-nitrilotriacetic acid affinity column. The column was washed with 20 mM Tris, pH 8.5, 200 mM NaCl, 5% glycerol, and 50 mM imidazole to remove nonspecifically bound proteins. Subsequently, Rv1601 was eluted from the column with 300 mM imidazole in the same buffer. The eluted protein was concentrated and further purified by size-exclusion chromatography using a HiLoad 16/600 Superdex 200 prep grade column (GE Healthcare) in 20 mM Tris, pH 8.5, and 200 mM NaCl buffer. The degree of purity of the IGPD (Rv1601) was examined by 10% SDS-PAGE.

### Biochemical assay

The activity of the enzyme was determined using a previously described stopped-assay protocol^22^ with minor modifications. Briefly, the reaction mixture consisted of 40 mM Triethanolamine (TEA) buffer pH 7.7, 50 mM MnCl_2_, 25 mM β-Mercaptoethanol, and 2 µg (1.16 µM) of the enzyme in a reaction volume of 75 μl. Initially, enzyme and substrate concentrations were optimized by bringing the output signal to the reading capacity of the instrument. Further, enzyme concentration was optimized for a linear range of initial velocity up to 60 s, maintaining the same substrate concentration. For the IC_50_ study, the reactions were carried out at 310 K using the fixed concentration of IGP (0.166 mM) and the different concentrations of triazole scaffold-based inhibitors (0.010 to 2.0 mM) in a serial 2-fold manner. The reactions were stopped by adding 250 μl of 1.43 M sodium hydroxide. Different time points were noted for the kinetics followed a gradation of 30 s. The reaction mixture was then incubated at 318 K for 20 min to convert the product imidazole acetol phosphate (IAP) into an enolized form, whose absorbance was read at 280 nm in a Shimadzu UV spectrophotometer against an appropriate blank. The extinction coefficient of IAP formed under these conditions was 5310 M^−1^ cm^−1^, as reported previously and this value was used for the calculation of kinetic parameters. Data points were plotted using Graph Pad Prism 6.0.

### x-ray crystal structures of enzyme/inhibitor complexes

To map the enzyme/inhibitor interactions, we carried out X-ray studies of enzyme/inhibitor complex. For this, IGPD crystals were soaked (grown using protocols as mentioned in our earlier study^12, 34^) in 0.5 mM of inhibitor solution for 10 min and subsequently diffraction data were collected and processed. The data collection and data processing statistics are presented in the Table 5. All the reported IGPD/inhibitor complex structures were refined in a similar manner. To begin with, each structure was subjected to 50 cycle of rigid body refinement followed by 100 cycles of maximum-likely-hood positional refinements The electron density maps for both 2*|F_o_|* - *|F_c_|* (at 1σ level) and |*F_o_|* - *|F_c_|* (at 3σ) were examined for the entire poly peptide chain and the model was corrected wherever necessary and subsequently water molecules were included into the model and positional and B-factor refinement were carried out. Based on our previous findings^12, 13^, and the feature that these inhibitors possess the scaffold similar to that of the imidazole ring of the substrate, each inhibitor is likely to bind in the enzyme’s active site. Examination of the 2*|F_o_|* - *|F_c_|* (at 1σ level) and |*F_o_|* - *|F_c_|* (at 3σ) confirmed the binding at the enzyme’s active site. Each inhibitor was modeled into the electron density and subsequently 100 cycles of positional refinement was carried out. The stereo chemical acceptability of all three enzyme/inhibitor complex structures were checked using Ramachandran plot analysis^35^ by PROCHECK^36^.

**Table 5:**
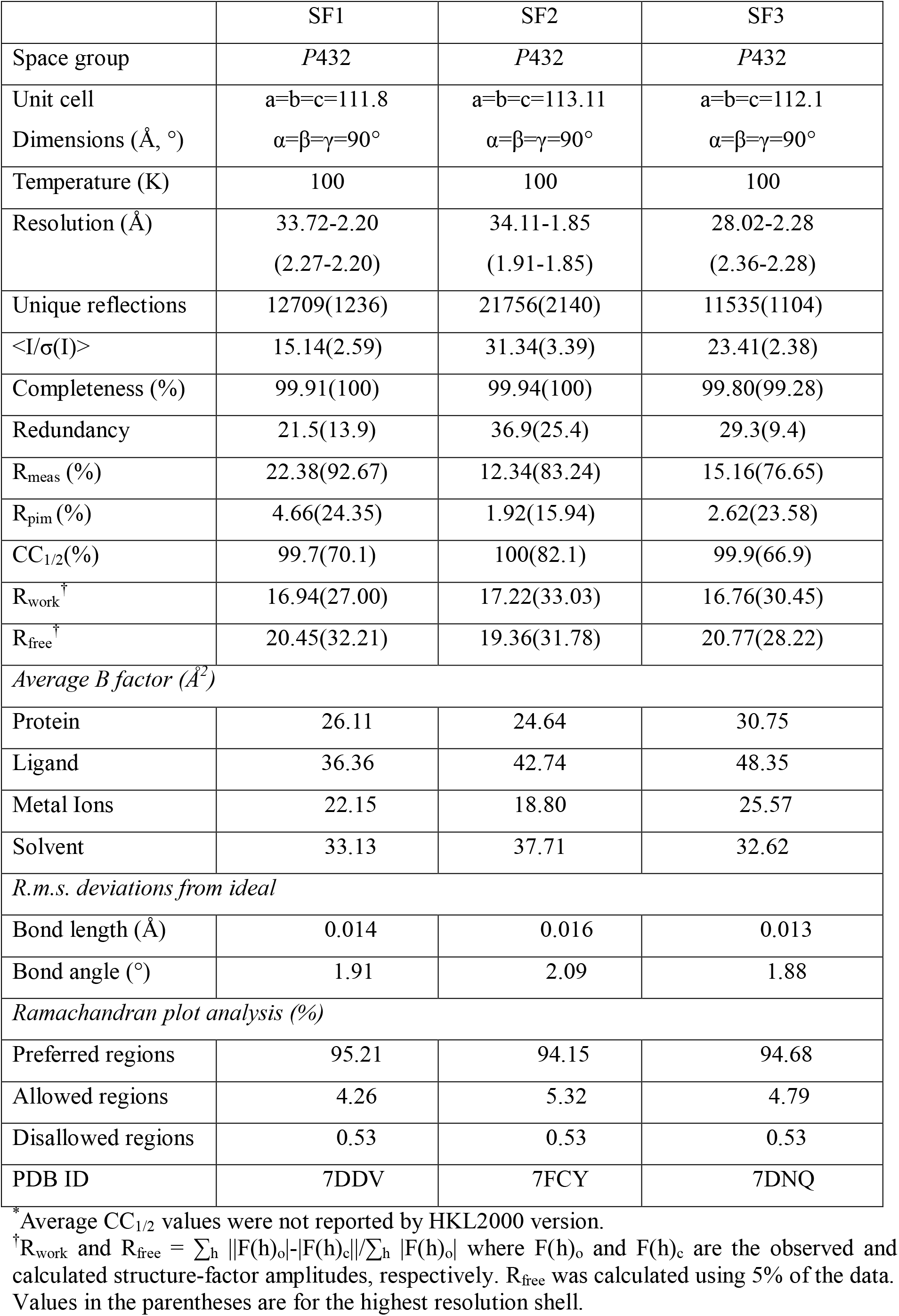
Data collection and refinement statistics.

### MIC determination

The antitubercular activity of the compound was tested by performing a Resazurin Assay (REMA). Briefly, *Mtb* H37Rv were grown to mid-logarithmic phase (A_600_ = 1-1.5) in Middlebrook 7H9 broth (Difco) supplemented with 10% OADC, 0.2% glycerol and 0.05% Tween 80 under aerobic conditions with shaking at 190 rpm. The culture was resuspended by passaging it 10–15 times through a 26.5-gauge needle and diluting it in growth medium to A_600_ = 0.001. All cultures were grown at 37°C in a shaker incubator. Drug susceptibility testing using resazurin was performed under aerobic conditions similar to that described for *Mtb* with minor modifications. The assay was performed in clear-bottomed, 96-well microplates (NUNC-Microwell, Thermo scientific). An initial stock solution of inhibitor was prepared in sterile deionized water, and subsequent 2-fold serial dilutions were prepared in 0.1 ml of 7H9-S/Dubos-S medium supplemented with 0.2% glycerol (without Tween 80) in the 96-well microplates. Approximately 10^4^ cells were added per well in a volume of 0.1 ml. Control wells containing [*Mtb* only and Medium + Compound (M+C)] were included in the plate setup. The plates were incubated at 37°C for 5 days. After that, 20 μl of 0.02% resazurin and 12.5 μl of 20% Tween 80 were added. The wells were observed after 24 and 48 h for a color change from blue to pink. Visual minimum inhibitory concentration (MIC) was defined as the lowest concentration of drug that prevented a color change.

### Macrophage infection

H37Rv bacterial strain was grown in Middlebrook 7H9 broth (Difco) supplemented with 10% OADC, 0.2% glycerol and 0.05% Tween-80 until the mid-log phase. The bacteria were harvested, washed with RPMI 1640 and re-suspended in the same media (Antibiotic Free). Bacteria were quantified by measuring the absorbance at a wavelength 600 nm. The Raw cells 264.7 (macrophages) were seeded in 6 well plates at 10^6^ cells/well and incubated for 3 h at 37°C. Macrophage monolayer was then washed with PBST and incubated with 2 ml infection media (RPMI 1640 medium supplemented with 10% FBS) containing 1×10^7^ cells of *Mtb* H37Rv (MOI =10) at 37°C under 5% CO_2_ for 3 h. Then the cells were washed twice with warm PBST to remove extracellular bacteria and supplemented with RPMI (without antibiotics) and a range of 0, 15, 30, 60, 120, 240 µM of Test compounds were added in respective wells. After 72 h of infection, media was aspirated from infected macrophage wells, and 500 µl of ice-cold sterile lysis solution (0.05% SDS, w/v in H_2_O) was added to each well. The lysates were further centrifuged at 1500 rpm for 10 min and transferred to new tube and serial dilutions were prepared. These dilutions were plated on 7H10 agar plates supplemented with OADC. The plates were kept in a CO_2_ incubator for 3–4 weeks and subsequently the colony forming units (CFU) were counted.

### Growth Kinetics of Mtb strains

The *Mtb* strain H37Rv was revived from glycerol stock and/or single colony picked from 7H10 agar medium and inoculated into 10 ml of 7H9 broth medium. Upon reaching mid-logarithmic phase (A_600_ ∼ 0.8), the cultures were diluted again to A_600_ ∼ 0.05. Cultures were then challenged with 1x MIC of inhibitors. Cultures were maintained for 21 days at 37^ᵒ^C and 100 r.p.m. A_600_ was determined at specific intervals. The growth kinetics was analyzed for three different compounds in three independently grown bacterial cultures (n=3).

### Cytotoxicity assay

#### Cell culture maintenance

RAW 264.7, HEK293 and THP-1 human monocytic cell line was obtained from ATCC. Cells were cultured in RPMI medium supplemented with 10% of heat-inactivated fetal bovine serum, 100 μM of penicillin and streptomycin, 50 μM neomycin and 200 mM of Glutamine in incubator at 37 °C under 5% CO_2_ atmosphere. They were passaged every 2–3 days to prevent the cell density from exceeding one million per ml. Mycoplasma contamination was not found in any of the cells tested. For each set of experiments, THP-1 cells were differentiated into macrophages by exposure to 10 ng/ml (16 nM) of PMA in a plate appropriate to the test for 24 h, while PMA induced activation is not required for RAW 264.7 and HEK 293 cells. The density of 10^4^ cells/ml was used for all tests.

#### In vitro Assay

Cells which were in the log phase of growth were harvested and determined by cell count. 10^4^ cells/well were used for the assay (suggested cell density). The optimum cell density may vary between cell types. Cells were seeded in 96 well plates and exposed to test compounds for 72 h to determine the cytotoxic effect of test compounds on cell growth. After 72 h alamarBlue was added at an amount of 10% of the total media volume in the well. Plates were incubated with alamarBlue for 4-8 hr. The optimum incubation time varied between cell types. The viability of the cells was measured after 72 h incubation at 37 °C, under 5% CO_2_, using the alamarBlue assay by measuring the absorbance at different wavelengths of 570 nm and 600 nm after required incubation. All Cytotoxicity experiments were performed in biological duplicates.

### Acute Toxicity Study

Healthy CD-1 female mice with weight 34±4 grams were used. The animals were housed individually in cages and maintained in a standard laboratory environment. Experimental protocols for the animal experiments were approved by the Institutional Animal Ethics Committee (IAEC) (IAEC#611/22) upon the recommendations and standards prescribed by the Committee for the Purpose of Control and Supervision of Experiments on Animals (CPCSEA), Government of India. Acute toxicity study was performed as per the up-and-down-procedure of Organization for Economic Cooperation and Development (OECD) guidelines 425. A limit dose of 2000 mg/kg body weight of compound was used involving three mice. Each mouse was treated with a single oral dose of 5, 50, 500, 2000 mg/kg body weight on Day 0. Animals were observed individually at least once during the first 30 min after dosing, periodically during the first 24 h and daily thereafter, for a total of 14 days for any clinical signs of toxicity or mortality. At the end of 14 day’s observation period, the animals were anaesthetized, and their blood samples were collected through retro orbital bleeding for biochemical and histopathological studies, respectively. For biochemical analysis, blood was centrifuged at 4000 rpm at 4°C for 10 min, serum was separated and Serum Glutamate Pyruvic Transaminase (SGPT), Serum Glutamate Oxaloacetic Transaminase (SGOT), creatinine, and, urea were estimated using a semi-automated Biochemical coralyzer. For histopathological studies, the animals were euthanized and the livers and kidneys were harvested and examined macroscopically. These organs were then preserved in 10% formalin made in PBS for histopathological examinations by standard techniques.

### Sample preparation for H&E Staining experiments

For this, the liver and kidney tissue samples were embedded into wax blocks. The wax blocks were subjected to microtome to generate sections, which were then stained with Hematoxylin and Eosin dyes at a commercial facility. The tissue sections were viewed in a bright field microscope at 40x magnification.

### Animal Infection Study

Animal experiments were performed with approval from Institutional Animal Ethics Committee (IAEC Approval No: IAEC/THSTI/196) as per guidelines laid by CPCSEA, Government of India. Briefly, bacterial suspensions of wild type *Mtb* H37Rv at a density of 10^8^ cells/ml was prepared in normal saline and female BALB/c (n=5 per group) were infected via aerosol route using Inhalation Exposure System (Glas-Col). Deposition of the bacteria as baseline infection was assessed by CFU at Day 1 post infection. Infection was allowed to progress through week 4 post infection at which animal groups (n=5) were administered orally with test compounds (mg/kg of animal body weight) – SF1 (100 mg/kg), SF2 (100 mg/kg), SF2 2X (200 mg/kg), SF3 (100 mg/kg), INH (10 mg/kg) and L-Histidine (100 mg/kg) for 5 days/week. Animals were euthanized at indicated time points post treatment, organs (lungs and/or spleens) were harvested and bacterial burden were determined by homogenization, serial dilution and CFU plating.

## Acknowledgments

The in-house X-ray diffraction facility, RIGAKU FR-E+ SuperBright microfocus rotating anode dual-wavelength (Cu and Cr) X-ray generator mounted with RAXIS IV^++^ detectors, was established with financial support from the DBT. BKB received projects (BT/PR29075/BRB/10/1699/2018, BT/PR40325/BTIS/137/1/2020) funding from the Department of Biotechnology (DBT) and Indian Council of Medical Research (Ref. no. 58/18/2015/-BMS) Government of India, India and from NII core. We sincerely thank Dr. Ravi Kant Pal for his help in crystal screening, data collection and data processing. Dr. Amit Kumar Pandey duly acknowledges Small Animal Facility and Infectious Disease Research Facility (IDRF), THSTI for Animal clearance study.

## Declaration

The authors declare that they have no competing interest.

## Notes

### Competing Interest Statement

The authors have declared no competing interest.

