## Supplementary File for "Chemical inhibition of histidine biosynthesis curtails *M. tuberculosis* infection"

[4]Present address: Department of Bioengineering, University of Illinois, Urbana Champaign

*Running title: disrupting histidine biosynthesis of *M. tuberculosis*

Keywords: *Mycobacterium tuberculosis*; Tuberculosis; Histidine biosynthesis; Imidazole glycerol phosphate dehydratase; Drug target; Inhibitor design

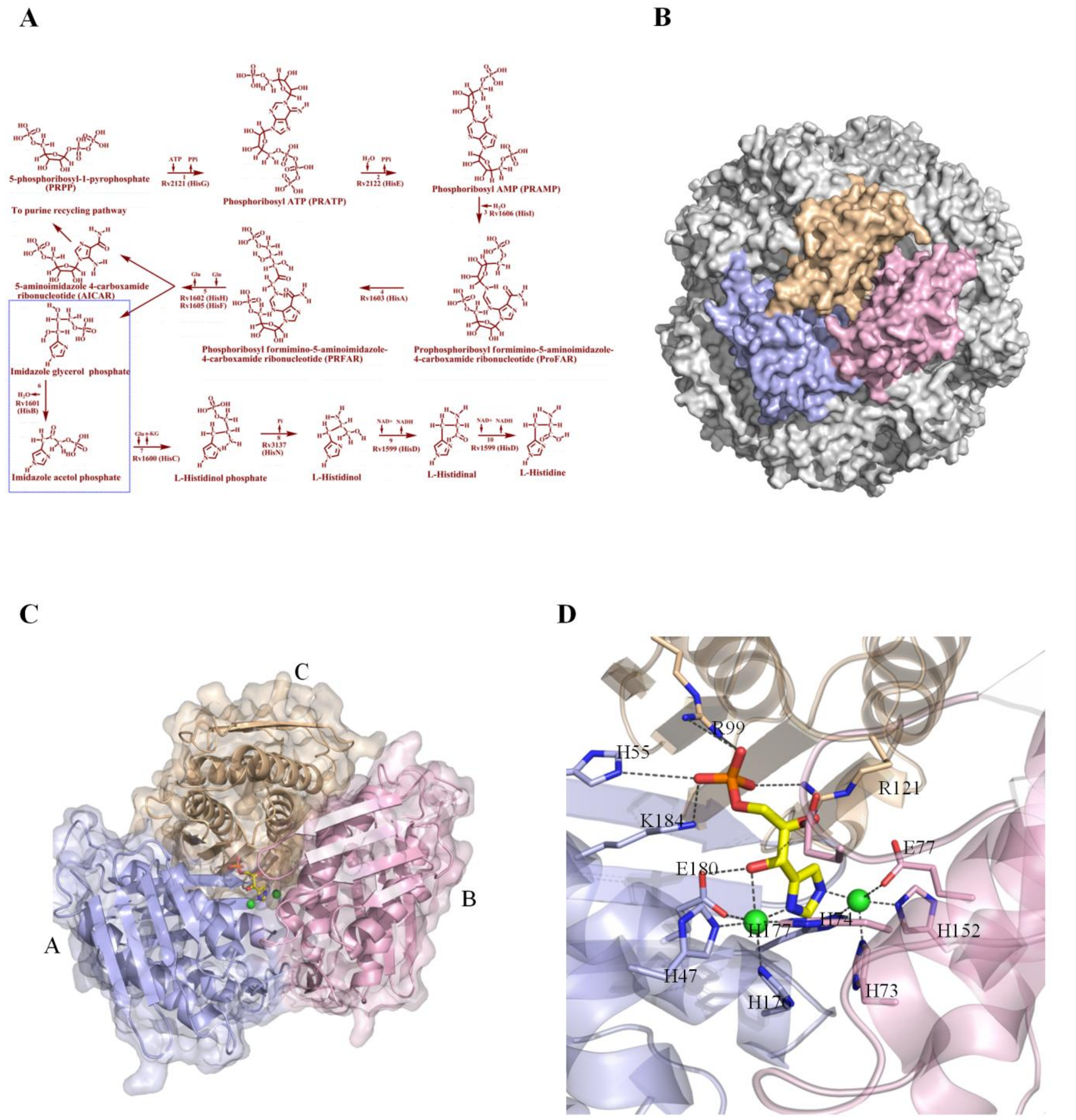

Figure S1. The involvement of HisB in the His pathway and its 3D structure and interactions with the substrate.

(A) Schematic representation of the reaction steps of histidine biosynthesis pathway[1]. The step that is catalyzed by HisB is defined by blue rectangle. (B) HisB functional unit, a 24-mer assembly shown as surface representation. Three colored monomers from one active site. (C) The active site composition of HisB[1]. (D) Interactions (shown in black dashes) of HisB with its substrate IGP[1,2]. The substrate is shown in stick representation with carbon atom in yellow color. The enzyme residues are shown in stick model, two active site $Mn^{2+}$ are shown in green spheres.

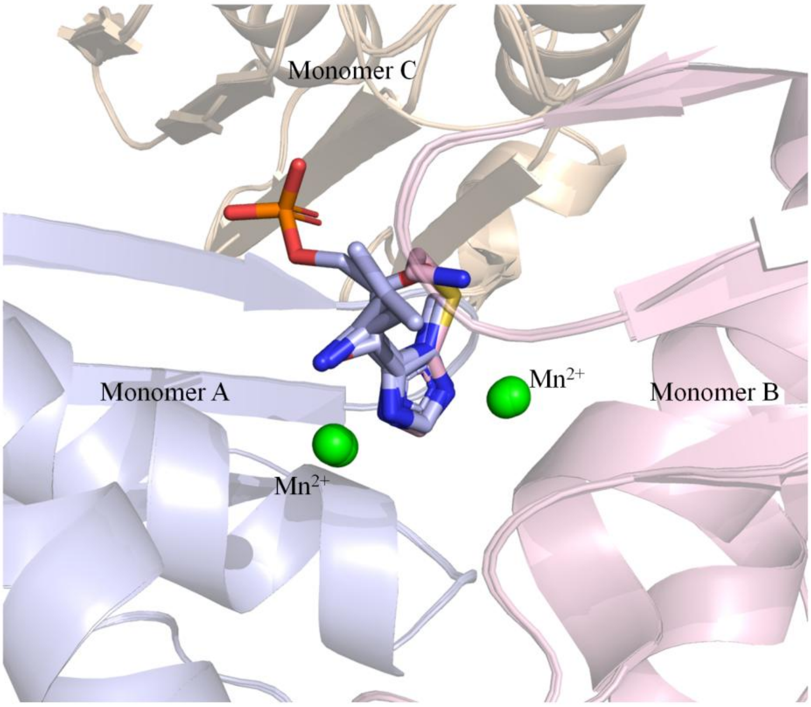

Figure S2. Superimposition of the HisB/IGP and HisB/SF1, SF2, SF3 structures.

IGP is shown in a stick model with carbon atoms in light blue, and the inhibitors are shown in a stick representation with carbon atoms also in light blue. Two $Mn^{2+}$ ions in the active site are shown as spheres in green color. The three monomers forming one active site unit are shown in the cartoon representation with monomer A (light blue), monomer B (light pink), and monomer C (wheat).

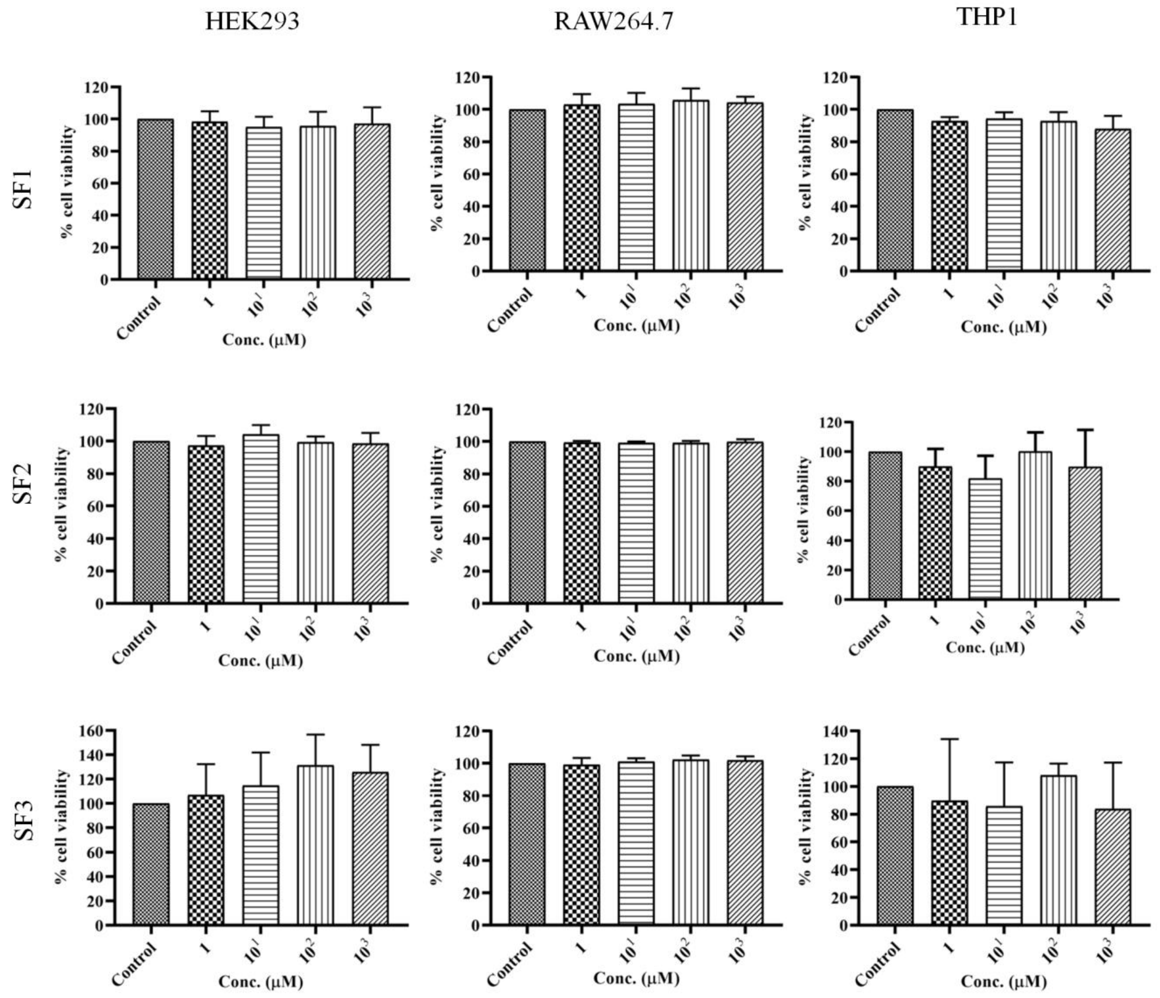

Figure S3. In vitro cytotoxicity study against different cell lines.

SF1, SF2, and SF3 are non-toxic to different cell lines. All compounds (SF1, SF2, and SF3) were tested for cytotoxicity on murine (RAW 264.7) and mammalian cells (THP1 and HEK293). No significant cytotoxicity was observed for these compounds.

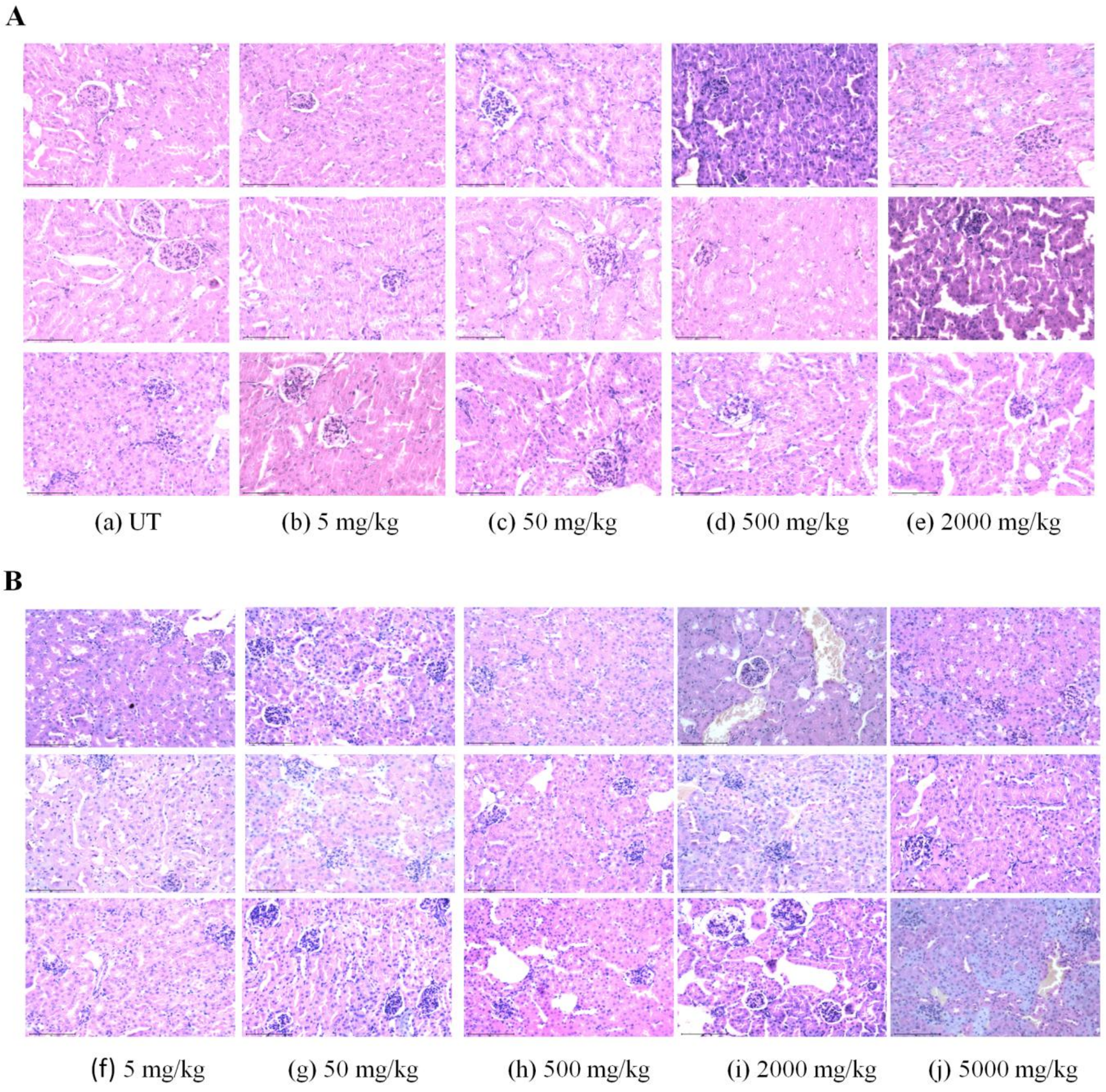

Figure S4. Histopathology of the kidney (hematoxylin and eosin staining, 40x_Scale bars: 100 µm).

(A) The effect of SF2 on kidney histology. Comparison of Untreated groups (UT) (a) vs. treated groups found to be normal with no pathological changes (b), (c), (d), and (e) indicates normal renal cortex, no leukocyte infiltration, no signs of edema, absence of necrotic foci, and normal tubular morphology at a dose of up to 2000 mg/kg body weight. (B) Effect of SF3 on kidney histology. Comparison of Untreated groups (UT) (a) vs. treated groups found to be normal with no pathological changes (f), (g), (h), (i), and (j) indicates normal renal cortex, no leukocyte infiltration, no signs of edema, absence of necrotic foci, and normal tubular morphology at a dose of up to 5000 mg/kg body weight.

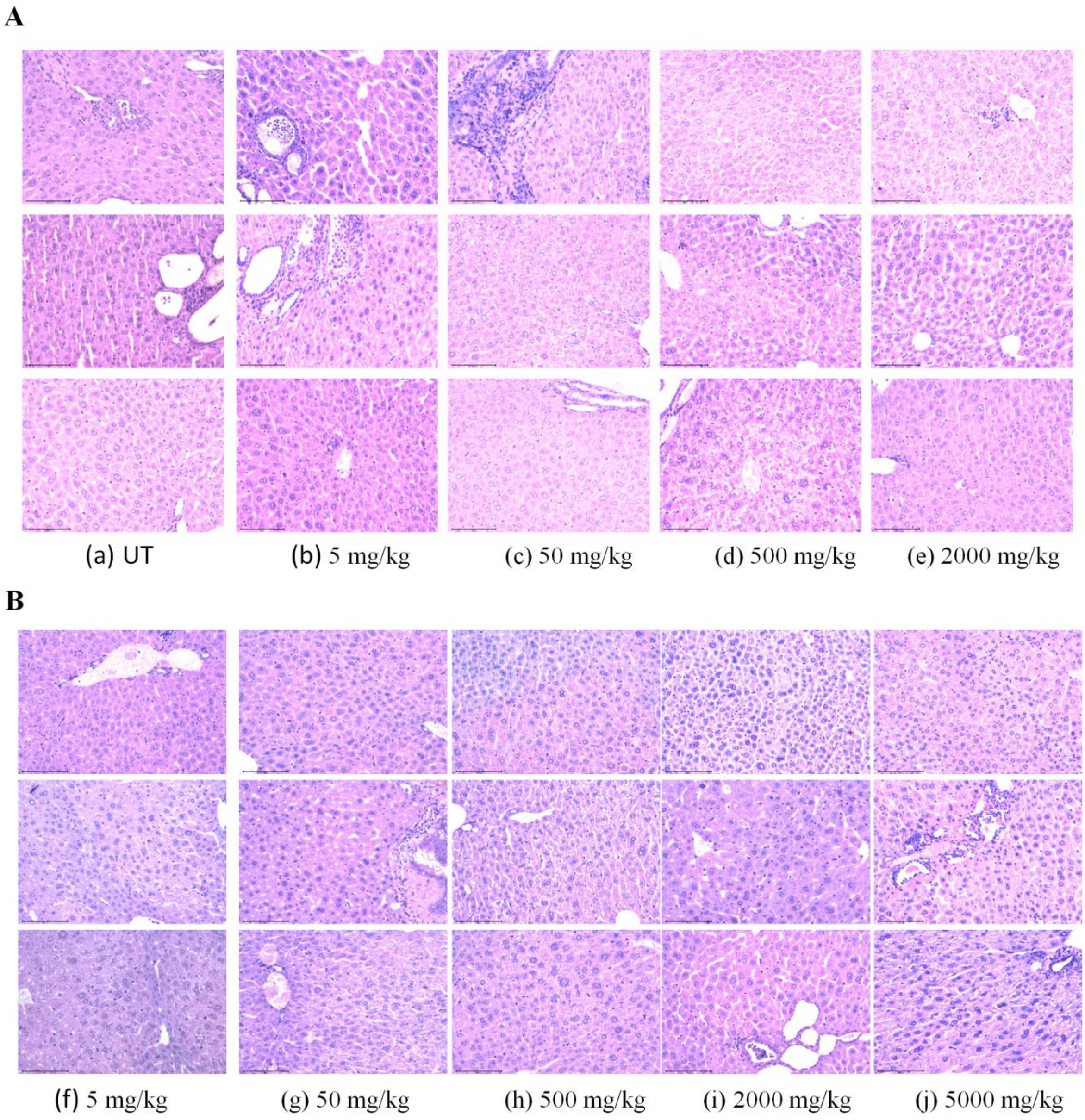

Figure S5. Histopathology of the liver (hematoxylin and eosin staining, 40x_Scale bar: 100 µm).

(A) Effect of SF2 on Liver histology. Comparison of Untreated groups (UT) (a) vs. treated groups found to be normal with no pathological changes (b), (c), (d), and (e) indicates normal Kupffer cells and no signs of granuloma, necrosis or infiltrations of degenerative cells at a dose of up to 2000 mg/kg body weight. (B) Effect of SF3 on Liver histology. Comparison of Untreated groups (UT) (a) vs. treated groups found to be normal with no pathological changes (f), (g), (h), (i), and (j), indicates normal Kupffer cells and no signs of granuloma, necrosis or infiltrations of degenerative cells at a dose of up to 5000 mg/kg body weight.

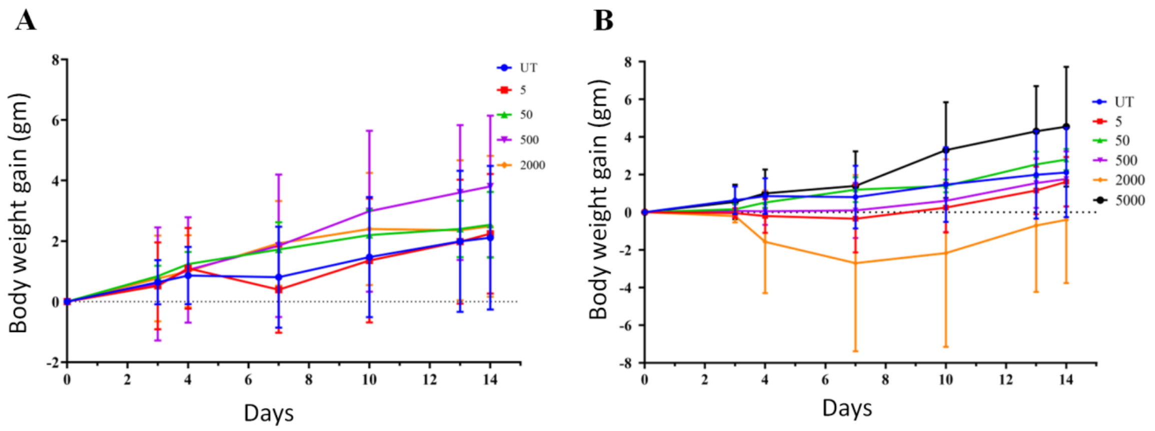

Figure S6. Acute toxicity study showing changes in mouse body weight.
(A) Effect of Compound SF2 at varying concentrations (5, 50, 500, 2000 mg/kg body weight) after a single dose administration on the body weight gain pattern of female mice monitored for 14 days. (B) Effect of Compound SF3 at varying concentrations (5, 50, 500, 2000, and 5000 mg/kg body weight) after a single dose administration on the body weight gain pattern of female mice monitored for 14 days. Data are shown as mean ± s.d (n = 5 for SF2 and SF3).

### Supplementary Tables

Table S1. Detail of in-house library compounds with their code, SMILE string, and Molecular formula.

| Code | SMILE string | Molecular Formula |
|---|---|---|
| SF1 | CC(C1=NC=NN1C)N | C5H10N4 |
| SF2 | CC(C)C(C1=NC=NN1C)N | C7H14N4 |
| SF3 | C1=NNC(=N1)SCCN | C4H8N4S |
| SF4 | C1=NNC(=N1)SCCCN=[N+]=[N-] | C5H8N6S |
| SF5 | C#CCNCCSC1=NC=NN1 | C7H10N4S |
| SF6 | CCNCC1=NC=NN1CC | C7H14N4 |
| SF7 | CCCNCC1=NC=NN1CC | C8H16N4 |
| SF8 | CC(C)NCC1=NC=NN1C | C7H14N4 |
| SF9 | CCNCC1=NC=NN1C | C6H12N4 |
| SF10 | CCCNCC1=NC=NN1C | C7H14N4 |
| SF11 | CCN1C(=NC=N1)CNC(C)C | C8H16N4 |
| SF12 | CCC1=NN(C(=N1)CNC(C)C)C | C9H18N4 |
| SF13 | C1C(CN1)SC2=NC=NN2 | C5H8N4S |

| | | |
|---|---|---|
| SF14 | CCCCN1C=NC(=N1)CN | C7H14N4 |
| SF15 | CC(C)N(C)CCNCC1=NC=NN1C | C10H21N5 |
| SF16 | CCC(CC)NCC1=NC=NN1C | C9H18N4 |
| SF17 | CCN1C(=NC=N1)CNCCC(C)C | C10H20N4 |
| SF18 | CCCN1C(=NC=N1)CNC | C7H14N4 |
| SF19 | CC(C)N1C(=NC=N1)CNCCN(C(C)C)C(C)C | C14H29N5 |
| SF20 | CC(C)NCC1=NC=NN1C(C)C | C9H18N4 |
| SF21 | CC(C)CN1C(=NC=N1)CNC(C)C | C10H20N4 |
| SF22 | CCCN1C(=NC=N1)CNC(C)C | C9H18N4 |
| SF23 | CCNCC1=NC=NN1C(C)C | C8H16N4 |
| SF24 | CCCN1C(=NC=N1)CNCC | C8H16N4 |
| SF25 | CCC(CC)NCC1=NC=NN1C(C)C | C11H22N4 |
| SF26 | CCCN1C(=NC=N1)CNC(CC)CC | C11H22N4 |
| SF27 | CCC(CC)NCC1=NC=NN1CC | C10H20N4 |
| SF28 | CCCNCC1=NC=NN1C(C)C | C9H18N4 |
| SF29 | CCCN1C(=NC=N1)CN(C)CCNC(C)C | C12H25N5 |
| SF30 | CCCN1C(=NC=N1)CN(CC)CCNC(C)C | C13H27N5 |
| SF31 | CCCNCCN(C)CC1=NC=NN1C(C)C | C12H25N5 |
| SF32 | CC(C)CCN1C(=NC=N1)CNC(C)C | C11H22N4 |
| SF33 | CC(C)CCN1C(=NC=N1)CNCC(C)C | C12H24N4 |
| SF34 | CC(C)CNCC1=NN(C=N1)C(C)C | C10H20N4 |
| SF35 | CC(C)CCN1C=NC(=N1)CNC(C)C | C11H22N4 |
| SF36 | CC(C)CCN1C=NC(=N1)CNC | C9H18N4 |
| SF37 | CCNCC1=NN(C=N1)CCC(C)C | C10H20N4 |
| SF38 | CCCNCC1=NN(C=N1)CCC(C)C | C11H22N4 |
| SF39 | CCC(CC)CN1C=NC(=N1)CNC(C)C | C12H24N4 |
| SF40 | CCC(CC)CN1C(=NC=N1)CNC(C)C | C12H24N4 |
| SF41 | CC(C)CCCN1C=NC(=N1)CNC(C)C | C12H24N4 |
| SF42 | CCCN(CC)CCNCC1=NC=NN1C(C)C | C13H27N5 |
| SF43 | CCCCNC(=NC)NCC1=NC=NN1C | C10H20N6 |
| SF44 | CCCN1C(=NC=N1)CNCCCN(C)C(C)C | C13H27N5 |
| SF45 | CCC(CC)CNCC1=NC=NN1C(C)C | C12H24N4 |
| SF46 | CCS(C)(C)C1=NC=NN1 | C6H13N3S |
| SF47 | CCC1=NC=NN1CCN | C6H12N4 |
| SF48 | C1=NNC(=N1)SCC=N | C4H6N4S |
| SF49 | CC(C)C1=NN(C=N1)C2CNC2 | C8H14N4 |
| SF50 | CN(C)CCSC1=NC=NN1 | C6H12N4S |

Table S2. General structures of compounds of in-house library with the respective codes.

| | |
|---|---|
| 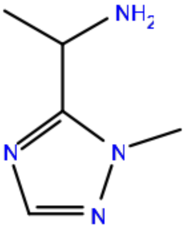 SF1 | 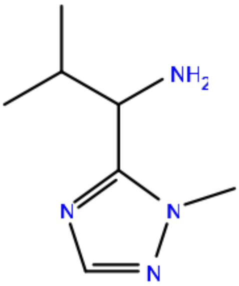 SF2 |
| 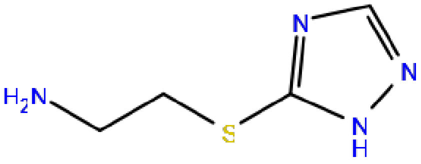 SF3 | 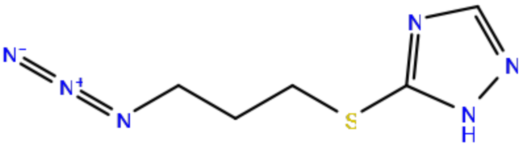 SF4 |
| 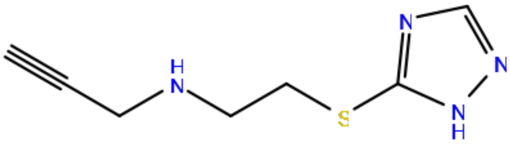 SF5 | 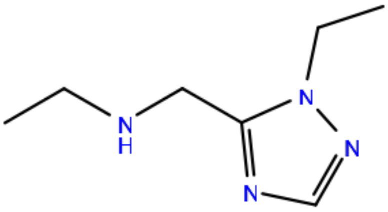 SF6 |
| 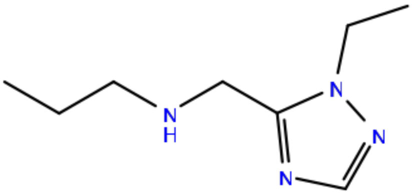 SF7 | 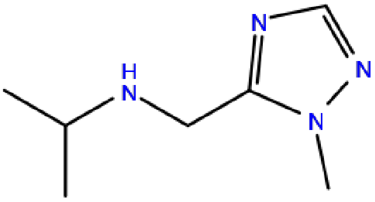 SF8 |
| 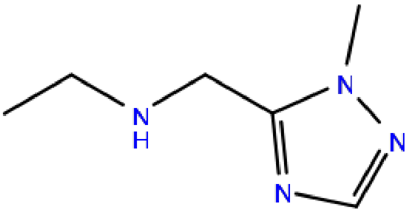 SF9 | 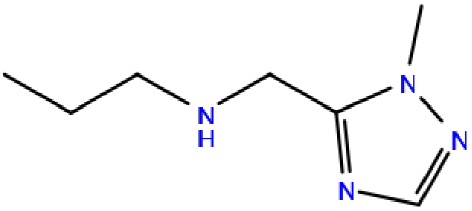 SF10 |
| 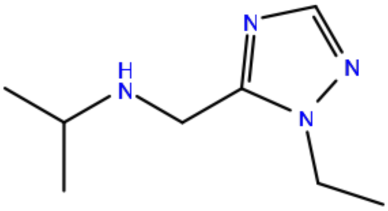 SF11 | 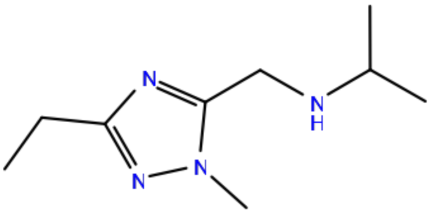 SF12 |

| | |
|---|---|
| 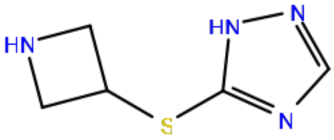 SF13 | 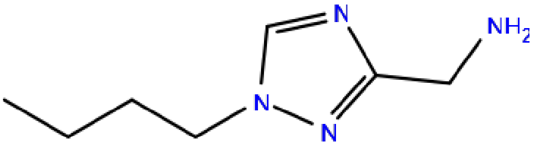 SF14 |
| 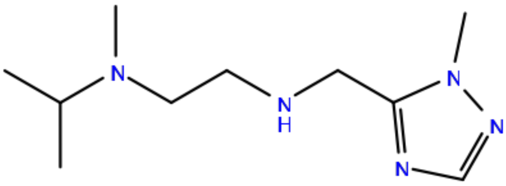 SF15 | 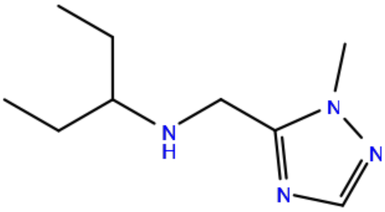 SF16 |
| 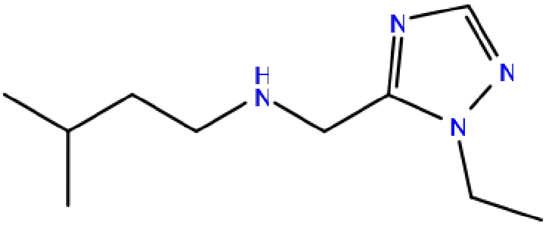 SF17 | 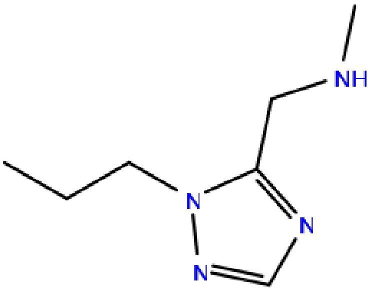 SF18 |
| 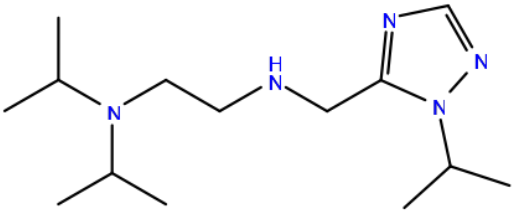 SF19 | 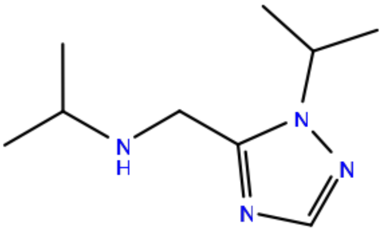 SF20 |
| 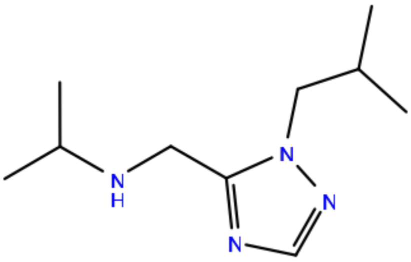 SF21 | 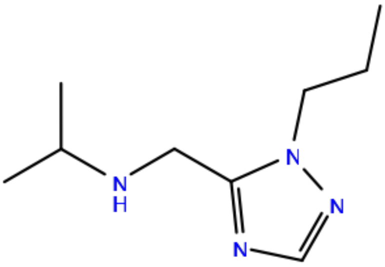 SF22 |
| 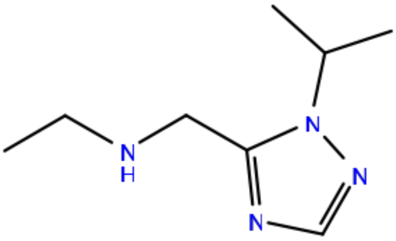 SF23 | 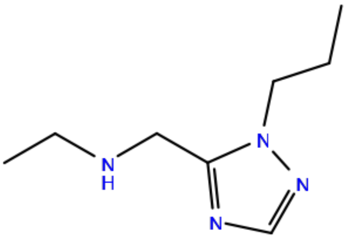 SF24 |

| | |
|---|---|
| 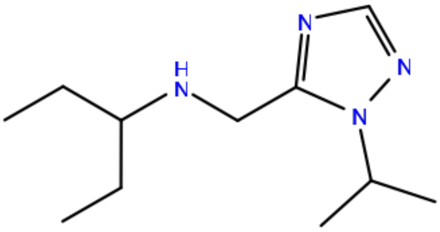<br>SF25 | 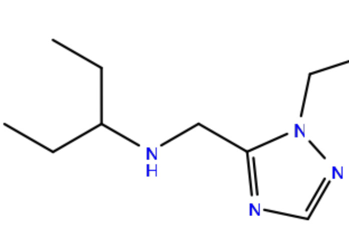<br>SF26 |
| 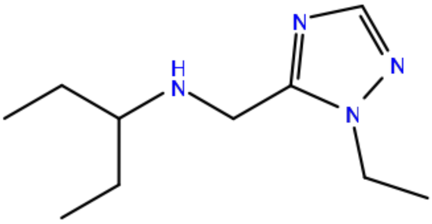<br>SF27 | 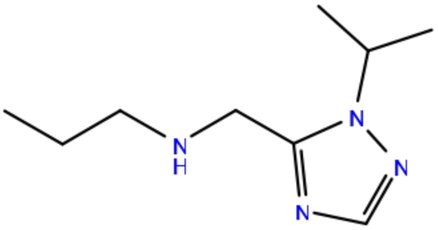<br>SF28 |
| 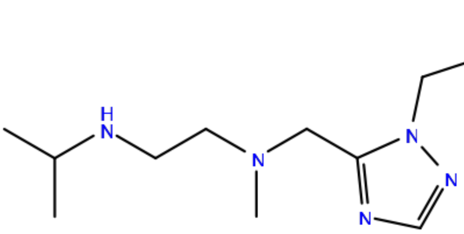<br>SF29 | 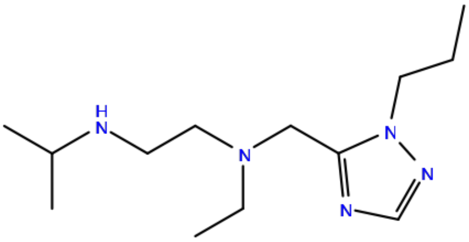<br>SF30 |
| 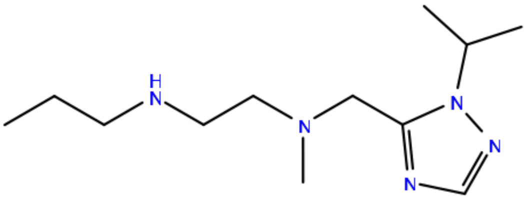<br>SF31 | 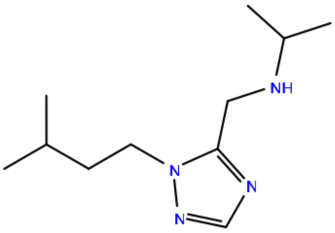<br>SF32 |
| 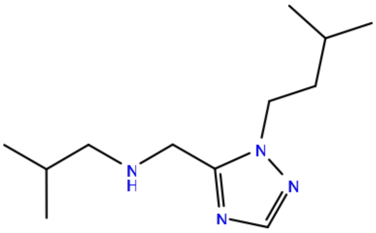<br>SF33 | 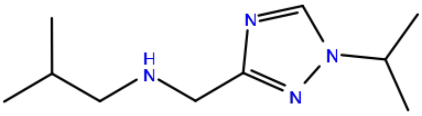<br>SF34 |
| 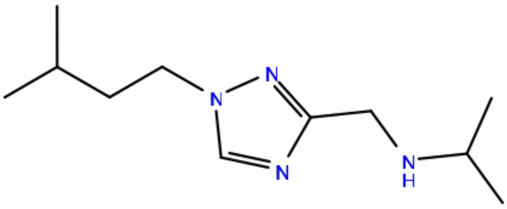<br>SF35 | 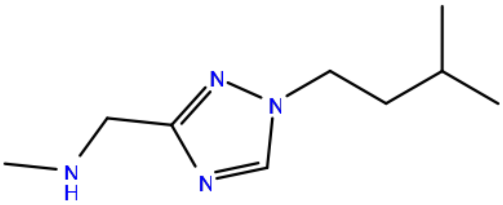<br>SF36 |

| | |
|---|---|
| 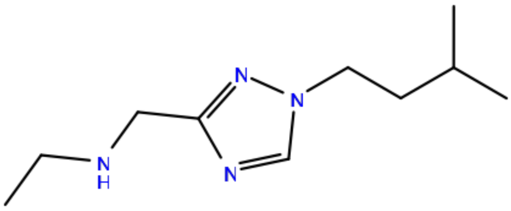<br>SF37 | 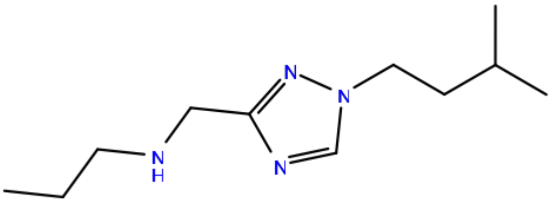<br>SF38 |
| 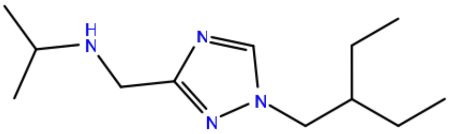<br>SF39 | 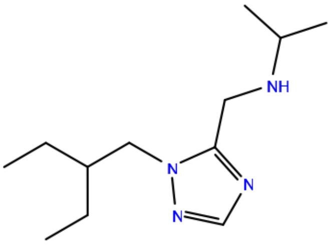<br>SF40 |
| 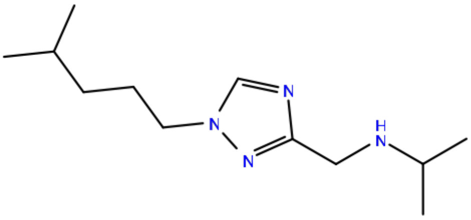<br>SF41 | 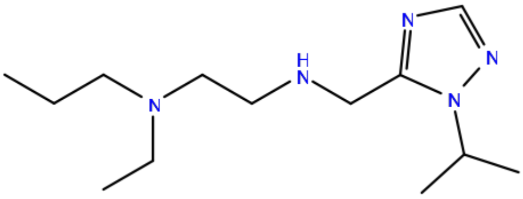<br>SF42 |
| 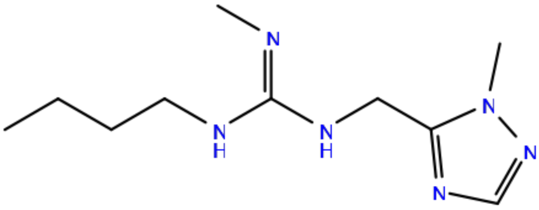<br>SF43 | 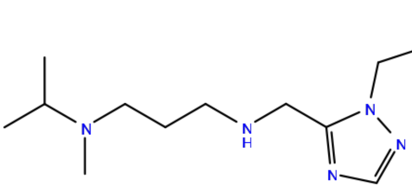<br>SF44 |
| 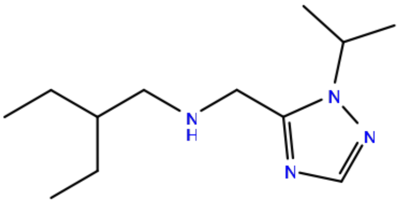<br>SF45 | 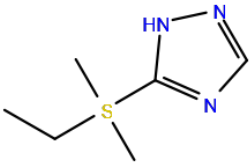<br>SF46 |
| 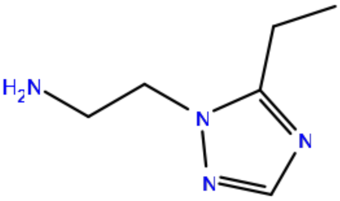<br>SF47 | 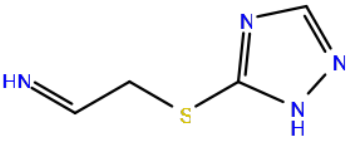<br>SF48 |

| | |
|---|---|
| 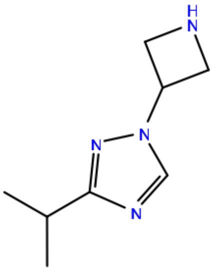 SF49 | 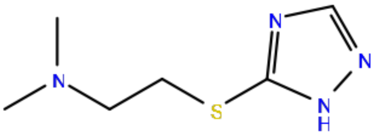 SF50 |
